# Coexistence of Cue-specific and Cue-independent Spatial Representations for Landmarks and Self-motion Cues in Human Retrosplenial Cortex

**DOI:** 10.1101/2022.05.16.491990

**Authors:** Xiaoli Chen, Ziwei Wei, Thomas Wolbers

## Abstract

Landmark-based and self-motion-based navigation are two fundamental forms of spatial navigation, which involve distinct cognitive mechanisms. A critical question is whether these two navigation modes invoke common or distinct spatial representations for a given environment in the brain. While a number of electrophysiological studies in non-human animals have investigated this question but yielded inconsistent results, it still awaits rigorous investigation in humans. In the current study, we combined ultra-high field fMRI at 7T and desktop virtual reality with state-of-the-art fMRI data analysis techniques. Using a novel linear track navigation task, we dissociated the use of landmarks and self-motion cues, so that participants used different spatial cues to encode and retrieve the same set of spatial locations. Focusing on the retrosplenial cortex (RSC) and the hippocampus, we observed that RSC contained both cue-specific and cue-independent spatial representations, which were driven by objective location (where the participant was actually located) and subjective location (the participant’s self-reported location), respectively. The hippocampus showed strong functional coupling with RSC and exhibited a similar spatial coding scheme, but with reduced effect sizes. Taken together, the current study demonstrated for the first time concurrent cue-specific and cue-independent spatial representations in RSC in the same spatial context, suggesting that this area might transform cue-specific spatial inputs into coherent cue-independent spatial representations to guide navigation behavior.

## INTRODCTION

The ability to localize and orient oneself as one navigates an environment is crucial for the survival of humans and non-human animals. Visual landmarks – salient objects in the environment – and self-motion cues represent two major and distinct types of cues used in spatial navigation. Landmark-based navigation is inherently discrete, in that a landmark can immediately inform about one’s whereabouts. On the contrary, self-motion cues are generated by one’s own movement and include body-based cues (e.g., vestibular feedback, proprioceptive cues, and motor efference copies) and optic flow. Navigation with self-motion cues alone is termed path integration, as one needs to infer self-position through continuous integration of self-motion inputs during locomotion.

Given the considerable body of evidence that landmark-based navigation and path integration recruit relatively independent cognitive ^1, 2^ and neural processes ^3, 4^, a critical question is whether these two navigation modes invoke common or distinct spatial representations in the brain. On the one hand, because landmarks and self-motion cues represent different sensory inputs, they may invoke separate neural representations of space. On the other hand, both cues typically denote the same space, hence spatial knowledge acquired from different cues should be integrated to generate a coherent representation that can guide navigation behavior. Deciding between these alternatives is fundamental to understanding the nature of cognitive maps, because it would provide important insights into an overarching question in spatial navigation – how different sources of spatial information are integrated to form a coherent cognitive map in the brain ^5–7^.

In non-human animals, cue-specific vs. cue-independent neural representations for the same environment have been examined intensively in the retrosplenial cortex (RSC) and hippocampus. For example, in bats, Geva-Sagiv and colleagues (2016) found that alternation between visual information and echolocation caused reorganization of hippocampal place fields within the same environment (i.e., remapping) ^9^, indicating that the hippocampus created cue-specific spatial maps even in the same environment. Studies in rodents usually manipulated the availability of visual information by switching a light on and off, but the results are mixed as to whether this manipulation would induce hippocampal remapping ^10–12^. Recently, Radvansky and colleagues (2021) showed that whether a common map or distinct maps were recruited for visual and odor cues depended on the behavioral relevance of these cues, i.e., whether different cue types were congruent or incongruent in defining a reward location ^13^. In RSC, researchers have also observed neurons exhibiting position-selective firing ^14, 15^; these place-cell-like cells maintained the same firing patterns when the environment was illuminated vs. dark ^14^, suggesting cue-independent spatial representations.

In humans, the question of cue-independent vs. cue-specific spatial maps has rarely been investigated. One notable exception is an fMRI study by Huffman & Ekstrom (2019) who varied the degree of body-based self-motion cues in different virtual reality environments ^16^. This study yielded preliminary evidence for cue-independent spatial representations in a large-scale brain network, as well as in RSC and the hippocampus. However, it is currently unknown whether the neural representations for a given environment are independent of or specific to the cue type used to encode and retrieve spatial locations.

To address this critical question, we employed ultra-high resolution fMRI at 7T, desktop virtual reality, and a mnemonic spatial navigation task to investigate whether cue-specific vs. cue-independent spatial representations are invoked by landmarks and self-motion cues. Specifically, we designed a spatial localization task in which participants encoded and retrieved the same set of four locations on a linear track, using either landmarks or self-motion cues alone; in other words, the two cue types were fully dissociated in the same spatial context (Figure 1a&b). We focused on RSC and the hippocampus, which have been investigated intensively in non-human animal studies on cue-specificity of spatial maps. We investigated spatial distance coding by exploiting two different types of fMRI effects well-suited to indexing neural representations of spatial relations – fMRI adaptation (fMRIa) and multi-voxel pattern similarity (MVPS). fMRIa and MVPS have been proposed to interrogate different aspects of the neuronal processing ^17^. Therefore, by investigating both effects, we aimed to obtain a more complete understanding of the neural mechanisms underlying spatial navigation in multi-information environments.

**Figure 1.**
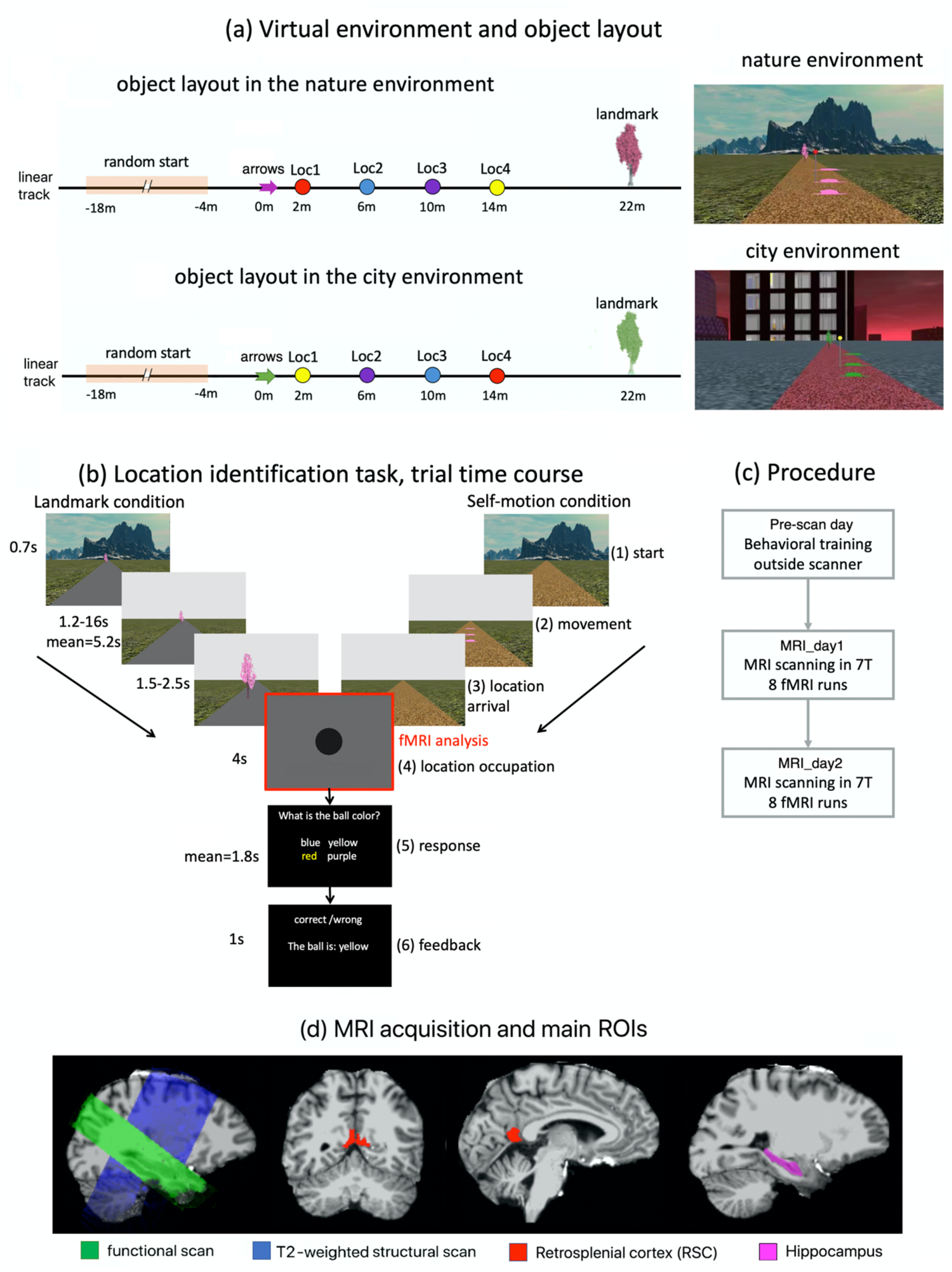
Environmental setup, navigation task, and MRI acquisition. **(a)** There were two different virtual environments (left): nature (upper panel) and city (lower panel). The two environments shared the same object layout on the linear track (left). There were arrows, four differently colored balls on poles, and a tree on the linear track. The four balls were positioned at the four test locations, i.e., Loc1, Loc2, Loc3, and Loc4. To improve visibility, we used three identical arrows positioned above the ground to denote the same spatial position, meaning that the arrows vertically projected to the same position on the ground and only differed in height. The arrows, the tree, and the floor texture of the linear track had the same physical appearances but in different colors in the two environments. The four balls positioned at the test locations were the same but reversed in order in the two environments. The floor texture outside of the linear track also differed between the two environments. Displayed on the right are snapshots of the two environments, with the background environment, the linear track, the tree, the arrows, and the ball positioned closest to the arrows. **(b)** The time course of the location identification task. Here, the trial is depicted in the nature environment, which was exactly the same in the city environment. Each trial had six phases. In phase 1 ‘start’, the participant was positioned at the starting location, which was randomized trial by trial based on a uniform distribution [−18 m, −4 m] (see Figure 1a, right). In phase 2 ‘movement’, the participant was passively transported to one of the four test locations. In phase 3, after arriving at the test location, the participant’s first-person perspective was smoothly turned down to vertically face the ground. In phase 4 ‘location occupation’, the participant’s perspective was fixed at the ground for four seconds. In phase 5 ‘response’, participant was required to identify the color of the ball positioned at that location within 20 second. In phase 6 ‘feedback’, feedback was provided, telling the participant whether the response was accurate, and, if incorrect, what the correct answer was. Note that the balls remained invisible throughout the trial, so that participants needed to recall from memory the color of the ball associated with the test location. In the landmark condition, the arrows were invisible, the tree was displayed, and the floor of linear track remained blank. In the self-motion condition, the arrows were displayed, the tree was invisible, and the texture of the linear track was displayed. In both conditions, the background environment only appeared briefly at the beginning of the trial (= 0.7s), and disappeared once the passive movement started. The fMRI analyses focused on the 4-second location occupation period (i.e., phase 4), when the visual inputs were the same for both cue conditions. **(c)** Participants were familiarized with the virtual environments and trained in the location identification task on the first day (Pre-scan day). On the following two days (MRI_day1 & MRI_day2), they completed the location identification task while undergoing MRI scanning in the 7T scanner. **(d)** MRI scanning and regions of interest. For an exemplary participant, the functional scan (in green), the T2-weighted structural scan (in blue), the anatomical mask of retrosplenial cortex (RSC; in red), and the anatomical mask of hippocampus (in violet) were overlaid on the brain extracted from the T1-weighted structural scan. For full details of the virtual environments and the experimental tasks, see STAR Methods and the video.

To preview, we found the most pronounced effects in RSC, which displayed both cue-specific and cue-independent spatial coding for landmarks and self-motion cues. Cue-specific coding was revealed by fMRIa and driven by objective location (i.e., the stimulus input, where the participant was actually located), whereas cue-independent coding was revealed by MVPS and driven by subjective location (i.e., the response output, where the participant thought they were located), indicating that RSC might transform cue-specific spatial inputs into abstract cue-independent spatial representations. The hippocampus exhibited a spatial coding scheme similar to that of RSC, but with reduced effect sizes. Taken together, the current study demonstrates for the first time the coexistence of cue-specific and cue-independent spatial representations in the human RSC.

## RESULTS

Twenty young healthy participants took part in the experiment. There were two different environments that shared the same object layout (Figure 1a). Participants needed to memorize the positions of four test locations that were evenly spaced along a linear track in a desktop virtual reality setup. Four balls of different colors were positioned at the four test locations. Participants performed a location identification task through the first-person perspective while undergoing MRI scanning at 7T on two consecutive days (Figure 1b, STAR Methods). In each trial, the participant was passively moved to a test location, and needed to identify the test location by recalling the color of the ball positioned at the location. The ball remained invisible throughout the trial. The arrows and the tree were positioned at the two ends of the ball object layout in the linear track, with the arrows closer to the starting position of the passive movement. We dissociated the use of landmark cues (a tree) and self-motion cues (optic flow elicited by the ground texture) in the task, so that on a given trial subjects could use only one cue type to encode and retrieve the test locations in the same environment. In the landmark condition, the tree served as the anchoring point, because it was the only spatial cue available and was informative of the participant’s self-position. In the self-motion condition, the position of the arrows served as the anchoring point, because once the participant had moved past the arrows, there were no further landmarks in sight, forcing the participant to estimate the travelled distance relative to the arrows by continuously integrating optic flow inputs.

### Behavioral evidence for a dissociation of landmarks and self-motion cues

Behavioral results are summarized in Figure 2. We submitted behavioral accuracy score to a repeated-measures ANOVA, with cue type, test location, scanning day, and environment as independent variables. This analysis revealed main effects of cue type (F(1,19) = 10.552, p = 0.004, η_p_^2^ = 0.357) and location (F(3,57) = 9.170, p < 0.001, η_p_^2^ = 0.326), which were qualified by a significant interaction between the two factors (F(3,57) = 25.051, p < 0.001, η_p_^2^ = 0.569) (Figure 2a): in the landmark condition, behavioral accuracy increased as the test location got closer to the tree (i.e., the anchoring point for landmark-based navigation), whereas in the self-motion condition, behavioral accuracy increased as the test location got closer to the arrows (i.e., the anchoring point for path integration). Accordingly, the interaction between cue type and the linear trend of test location was significant (t(57) = 8.487, p < 0.001). A closer look revealed that the linear trend of test location was significant in both the landmark condition (t(112) = 3.020, p = 0.003) and the self-motion condition (t(112) = 9.798, p < 0.001). No effects involving scanning day or environment were significant (ps > 0.3).

**Figure 2.**
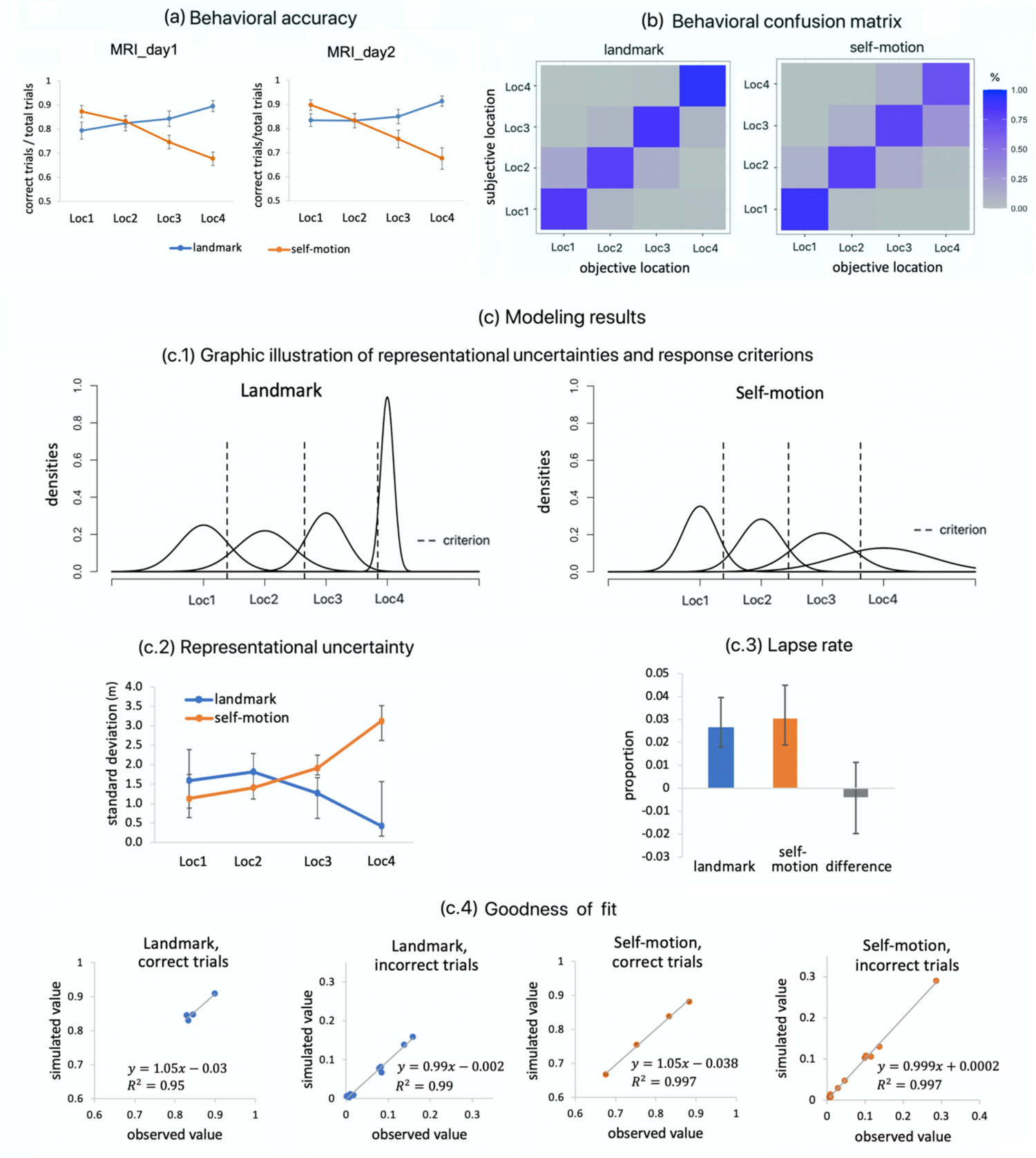
Behavioral results. (a) Behavioral accuracy is plotted as a function of test location and cue type in each scanning day. Error bars represent ± S.E. (b) Behavioral confusion matrix. Columns represent the objective locations (i.e., where the participant was actually located), whereas rows represent the subjective locations (i.e., where the participant reported he/she was located). Each cell represents the proportion of trials falling into the category. (c) Disentangling representational precision and response bias via modeling. (c.1) Graphic illustration of the estimated underlying representations and response criteria. (c.2) Estimated representational uncertainty (i.e., standard deviations of the Gaussian distributions in (c.1)) is plotted as a function of location and cue type. Error bars represent 95% confidence intervals obtained through a bootstrapping procedure. We further found that the interaction effect between cue type and the linear trend of test location on representational uncertainty was significant (i.e., the 95% confidence interval of the interaction effect did not contain zero). (c.3) Lapse rate was significantly higher than zero in both the landmark condition and the self-motion condition (i.e., the 95% confidence intervals did not contain zero), indicating that participants failed to pay adequate attention to the task occasionally. The difference in lapse rate was not significant between the two cue types. Error bars represent 95% confidence intervals obtained through a bootstrapping procedure. (c.4) Goodness-of-fit of the model. Regarding the behavioral confusion matrices, the observed values in the observed matrices are plotted against the simulated values generated by the model using the optimal values of the parameters as depicted in (c.1), separately for landmarks and self-motion cues, and separately for correct and incorrect trials. The linear regression line, R^2^ (i.e., goodness-of-fit), and the regression equation are displayed in each scatterplot. Additionally, goodness-of-fit remained at a very high level when all data points were analyzed together (R^2^ = 0.9997). See more details of the analyses in STAR Methods.

Since behavioral accuracy is jointly determined by representational precision and response bias, we tested whether using different navigational cues affected the underlying cognitive representations of the test locations. First, we aggregated data across participants and computed a group-level behavioral confusion matrix to characterize how participants confused the test locations (e.g., choosing location 1 as the response while actually occupying location 2). As can be seen in Figure 2b, mistakes mostly occurred between adjacent locations (e.g., Loc1 & Loc2), but rarely between locations that were farther apart (e.g., Loc1 & Loc4). Next, the group-level behavioral confusion matrices were submitted to an extension of signal detection theory, which we developed to accommodate tasks with more than two choices (Figure 2c; STAR Methods). The results showed that this model fitted the data very well (Rs > 0.94; Figure 2c.4). Furthermore, as shown in Figure 2c.1 and Figure 2c.2, in both cue conditions, the standard deviation of the underlying representation for the test location (i.e., the inverse of precision) decreased as the test location became closer to the respective anchoring points (i.e., the tree in the landmark condition and the arrows in the self-motion condition), which corresponds to the behavioral accuracy results (Figure 2a).

In summary, the primary behavioral finding was the differential performance profiles across test locations in the two cue conditions. Importantly, this finding was not confounded by possible response biases, because the underlying representational precision was influenced by test location and cue type in the same manner. Taken together, the behavioral results indicated that our cue dissociation manipulation was successful.

### fMRI results

fMRI analyses focused on the location occupation phase of the location identification task, when the camera was panned down to the ground to render visual inputs identical between the landmark condition and self-motion condition (Figure 1b). We examined whether BOLD signals in RSC contained information about allocentric distance-based spatial relations among the test locations. To this end, by adopting a continuous carry-over design ^18, 19^, we simultaneously investigated fMRI adaptation (fMRIa) and multi-voxel pattern similarity (MVPS), both of which have been used to evaluate neural representations of allocentric spatial relations between locations ^20^. fMRIa refers to the phenomenon that the BOLD signal is reduced if the current location is preceded by the same or a nearby location, and the degree of repetition suppression is proportional to the spatial proximity of the two locations ^3, 21, 22^. MVPS exploits the voxel-to-voxel distribution of brain activation that indexes neural representation, based on the rationale that spatial locations that are closer to each other should evoke more similar neural representations ^23, 24^. One hypothesis postulates that fMRIa and MVPS respectively interrogate the neuronal input stage, which is associated with stimulus input, and the neuronal output stage, which is associated with response output ^17^. This hypothesis has received empirical support (see discussion for details). Motivated by this hypothesis, we analyzed fMRIa and MVPS in terms of both stimulus input (e.g., objective location, where the participant was actually located) and response output (i.e., subjective location, where the participant reported he/she was located), and then corrected for multiple comparisons. fMRI analyses focused on the RSC and the hippocampus. For completeness, results for other areas in the medial temporal lobe (i.e., parahippocampal cortex, perirhinal cortex, and entorhinal subregions) are summarized in the supplemental information (Table S3).

### RSC showed fMRIa-based spatial distance coding for both landmarks and self-motion cues, which was driven by objective location

To examine whether RSC encoded spatial distance information in the form of fMRIa, we conducted univariate fMRIa analyses, in which we included parametric regressors that modeled modulatory effects of spatial distances between successively visited test locations in the first-level general linear models (STAR Methods, fMRIa-GLM1). The parametric regressors were defined either by objective or subjective locations. In the self-motion condition, the parametric regressors modeled spatial distances between successively visited test locations in a continuous manner by default, with four possible values of 0m, 4m, 8m, and 12m. Given a previous study showing that in retrosplenial regions fMRIa associated with landmark-defined locations only differentiated between same vs. different locations (but not between different spatial distances between different locations) ^22^, the parametric regressors in our landmark condition reflected whether the currently visited location was identical to (value = 0) or different from (value = 1) the preceding one. For each parametric regressor, beta estimates were averaged across all voxels in RSC.

We analyzed fMRIa averaged across the two environments and the two scanning days. We tested objective-location-based and subjective-location-based fMRIa for the two cue conditions against 0 using one-tailed simple t tests, with the familywise type I error controlled at 0.05 using the permutation-based Holm-Bonferroni method for the four individual t tests (STAR Methods). As shown in Figure 3a and Table 1, RSC showed significant objective-location-based fMRIa for landmarks (p_corrected_ = 0.005) and self-motion cues (p_corrected_ = 0.012). The subjective-location-based fMRIa was significant in the landmark condition (p_corrected_ = 0.010) but not in the self-motion condition (p_corrected_ = 0.061). Additional analyses showed that in the landmark condition, fMRIa was stronger on the 2^nd^ than the 1^st^ scanning day (p = 0.012): fMRIa was highly significantly on the 2^nd^ day (objective location, p_1-tailed,uncorrected_ = 0.0004, BF_10_= 80.494; subjective location, p_1-tailed,uncorrected_ = 0.002, BF_10_ = 18.596), but not significant on the 1^st^ day (ps_1-tailed,uncorrected_ > 0.45, BFs_10_ < 0.25). No significant influences of environment were observed (Table S1). Overall, these results suggest that fMRIa was associated more strongly with objective location than subjective location.

**Figure 3.**
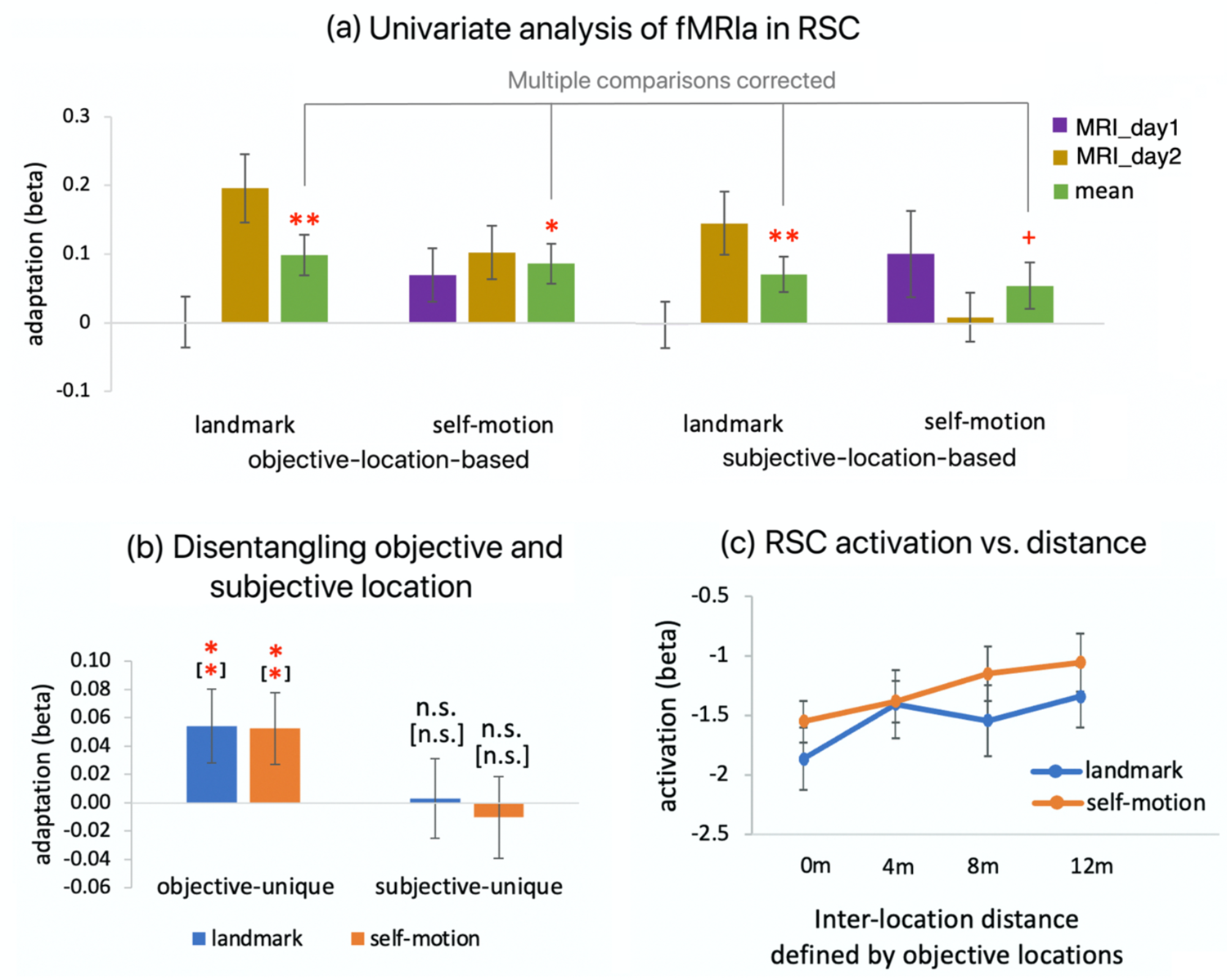
Univariate analysis of fMRIa in retrosplenial cortex. (a) Beta estimate of fMRIa is plotted as a function of location type (objective vs. subjective location), cue type (landmark vs. self-motion), and scanning day (MRI_day1 vs. MRI_day2). We conducted statistical tests on the mean fMRIa averaged across scanning days and environments (green bars). The displayed significance results were corrected for multiple comparisons of the four tests, using the permutation-based Holm-Bonferroni procedure. (b) Unique contributions of objective location (‘objective-unique’) and subjective location (‘subjective-unique’) to fMRIa. Results of the landmark condition and the self-motion condition are plotted separately. Significance levels displayed in the brackets refer to results when participants with behavioral accuracy > 90% were excluded from the analysis. (c) Beta estimate of RSC activation is plotted as a function of inter-location distance defined by objective locations for each cue type. See more details of the analysis in STAR Methods. n.s. denotes p_1-tailed_ > 0.1, * denotes p_1-tailed_ < 0.05, and ** denotes p_1-tailed_ < 0.01; + denotes p_1-tailed_ < 0.1.

**Table 1.**
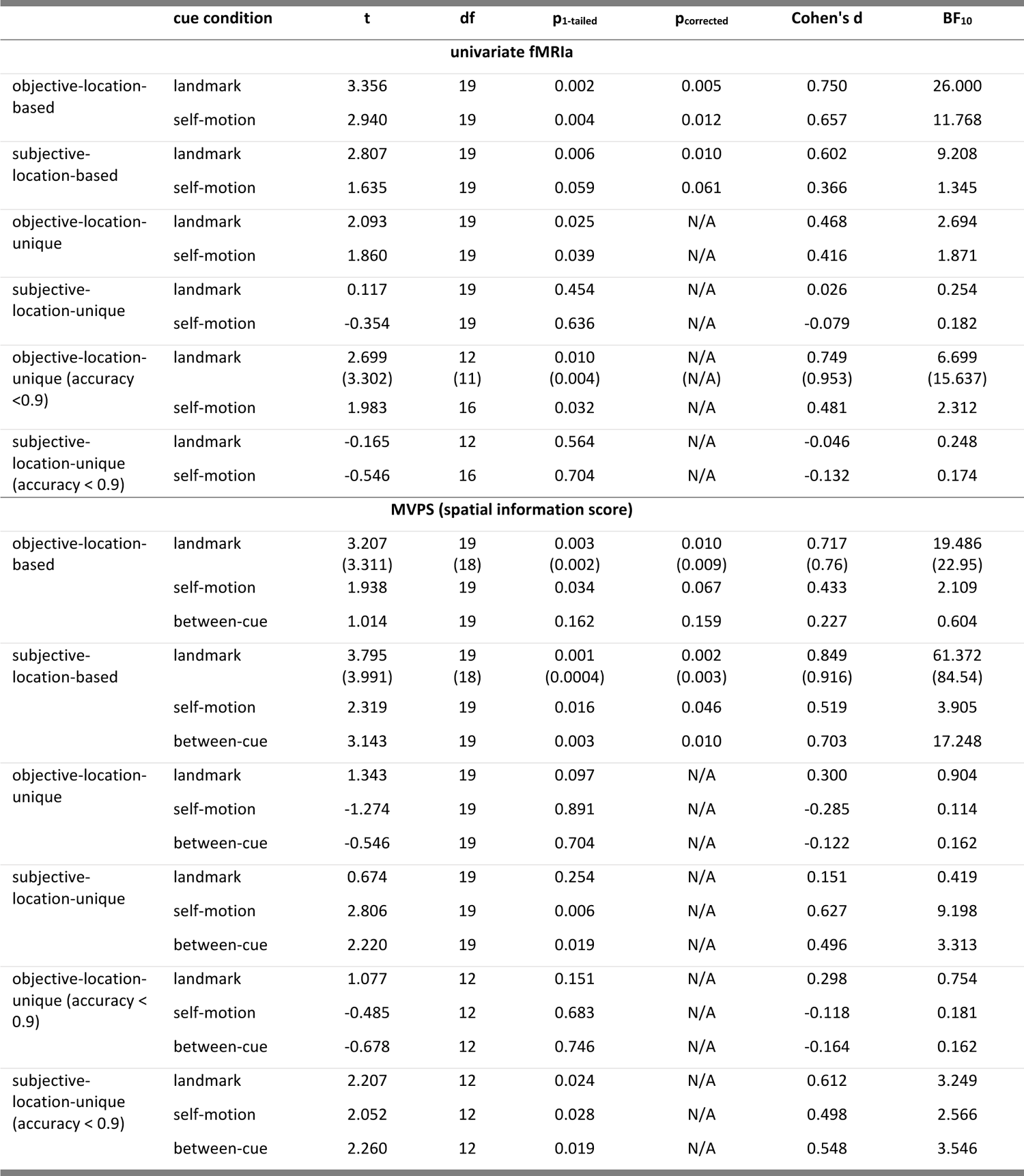
Statistical results for univariate fMRI adaptation and multi-voxel pattern similarity analyses. In correspondence to Figure 3a&b on fMRI adaptation (fMRIa) and Figure 5b&c on multi-voxel pattern similarity (MVPS). In paratheses are results with statistical outliers excluded from the analysis. df: degree of freedom; N/A: not applicable; BF_10_: Bayes factor, relative likelihood of the alternative hypothesis over the null hypothesis.

|  | cue condition | t | df | p<sub>1-tailed</sub> | p<sub>corrected</sub> | Cohen's d | BF<sub>10</sub> |
| --- | --- | --- | --- | --- | --- | --- | --- |
| univariate fMRIa |  |  |  |  |  |  |  |
| objective-location-based | landmark | 3.356 | 19 | 0.002 | 0.005 | 0.750 | 26.000 |
|  | self-motion | 2.940 | 19 | 0.004 | 0.012 | 0.657 | 11.768 |
| subjective-location-based | landmark | 2.807 | 19 | 0.006 | 0.010 | 0.602 | 9.208 |
|  | self-motion | 1.635 | 19 | 0.059 | 0.061 | 0.366 | 1.345 |
| objective-location-unique | landmark | 2.093 | 19 | 0.025 | N/A | 0.468 | 2.694 |
|  | self-motion | 1.860 | 19 | 0.039 | N/A | 0.416 | 1.871 |
| subjective-location-unique | landmark | 0.117 | 19 | 0.454 | N/A | 0.026 | 0.254 |
|  | self-motion | -0.354 | 19 | 0.636 | N/A | -0.079 | 0.182 |
| objective-location-unique (accuracy < 0.9) | landmark | 2.699<br>(3.302) | 12<br>(11) | 0.010<br>(0.004) | N/A<br>(N/A) | 0.749<br>(0.953) | 6.699<br>(15.637) |
|  | self-motion | 1.983 | 16 | 0.032 | N/A | 0.481 | 2.312 |
| subjective-location-unique (accuracy < 0.9) | landmark | -0.165 | 12 | 0.564 | N/A | -0.046 | 0.248 |
|  | self-motion | -0.546 | 16 | 0.704 | N/A | -0.132 | 0.174 |
| MVPS (spatial information score) |  |  |  |  |  |  |  |
| objective-location-based | landmark | 3.207<br>(3.311) | 19<br>(18) | 0.003<br>(0.002) | 0.010<br>(0.009) | 0.717<br>(0.76) | 19.486<br>(22.95) |
|  | self-motion | 1.938 | 19 | 0.034 | 0.067 | 0.433 | 2.109 |
|  | between-cue | 1.014 | 19 | 0.162 | 0.159 | 0.227 | 0.604 |
| subjective-location-based | landmark | 3.795<br>(3.991) | 19<br>(18) | 0.001<br>(0.0004) | 0.002<br>(0.003) | 0.849<br>(0.916) | 61.372<br>(84.54) |
|  | self-motion | 2.319 | 19 | 0.016 | 0.046 | 0.519 | 3.905 |
|  | between-cue | 3.143 | 19 | 0.003 | 0.010 | 0.703 | 17.248 |
| objective-location-unique | landmark | 1.343 | 19 | 0.097 | N/A | 0.300 | 0.904 |
|  | self-motion | -1.274 | 19 | 0.891 | N/A | -0.285 | 0.114 |
|  | between-cue | -0.546 | 19 | 0.704 | N/A | -0.122 | 0.162 |
| subjective-location-unique | landmark | 0.674 | 19 | 0.254 | N/A | 0.151 | 0.419 |
|  | self-motion | 2.806 | 19 | 0.006 | N/A | 0.627 | 9.198 |
|  | between-cue | 2.220 | 19 | 0.019 | N/A | 0.496 | 3.313 |
| objective-location-unique (accuracy < 0.9) | landmark | 1.077 | 12 | 0.151 | N/A | 0.298 | 0.754 |
|  | self-motion | -0.485 | 12 | 0.683 | N/A | -0.118 | 0.181 |
|  | between-cue | -0.678 | 12 | 0.746 | N/A | -0.164 | 0.162 |
| subjective-location-unique (accuracy < 0.9) | landmark | 2.207 | 12 | 0.024 | N/A | 0.612 | 3.249 |
|  | self-motion | 2.052 | 12 | 0.028 | N/A | 0.498 | 2.566 |
|  | between-cue | 2.260 | 12 | 0.019 | N/A | 0.548 | 3.546 |

To rigorously disentangle the contributions of objective vs. subjective location, we directly compared them by including parametric regressors for objective-location-defined and for subjective-location-defined spatial relations in the same first-level general linear model (STAR Methods, fMRIa-GLM2). As shown in Figure 3b and Table 1, the unique contribution of objective location was significant in both the landmark (p_1-tailed_ = 0.025) and the self-motion condition (p_1-tailed_ = 0.039). In contrast, the unique contribution of subjective location was not significant in either the landmark condition (p_1-tailed_ = 0.454) or the self-motion condition (p_1-tailed_ = 0.636). Given that high behavioral accuracy levels could cause unreliable beta estimates of the parametric regressors due to high correlations between objective-location-defined and subjective-location-defined spatial relations, we excluded participants with behavioral accuracy > 90% from the analysis; the pattern of results remained unchanged.

Additional analyses showed that these results could not be explained by potential differences in the relative detection power (DP_rel_) between the objective-location-defined and subjective-location-defined parametric regressors (STAR Methods). Although the objective-location sequences had significantly higher DP_rel_ for fMRIa than subjective-location sequences (p < 0.001), the magnitude of the difference was negligible (DP_rel_ = 64% vs. 63%).

Together, these results indicate that fMRIa was predominantly driven by objective rather than subjective location.

To visualize objective-location-based fMRIa (Figure 3c; STAR Methods, fMRIa-GLM3), in the self-motion condition, RSC activation increased in a linear manner as inter-location distance increased from 0m to 12m; in the landmark condition, RSC activation was higher at the non-zero inter-location distances relative to the zero inter-location distance (i.e., when two successively visited locations were the same), but remained at similar levels for different non-zero distances.

Finally, we conducted the voxel-wise analysis to investigate fMRIa in the entire-volume (Figure S1). In posterior cingulate areas (including RSC proper and the putative retrosplenial complex), fMRIa appeared to be stronger when based on objective location than subjective location in both cue conditions. This trend also existed in other brain regions, e.g., precuneus, calcarine, and angular gyrus.

To summarize, the results showed that RSC encoded spatial information for both cue types in the form of repetition suppression, which was mainly driven by objective rather than subjective location. For landmarks, the spatial coding mainly differentiated between same vs. different locations, whereas for self-motion cues the spatial coding differentiated different inter-location distances in a continuous manner.

### fMRIa-based distance coding was spatially distinct between cue types in RSC

To uncover whether or not the underlying fMRIa-based neural representations were distinct between the two cue types, we analyzed the similarity of the fMRIa patterns (Figure 4a; STAR Methods, fMRIa-GLM1), i.e., we determined whether voxels showing higher fMRIa for one cue also showed higher fMRIa for the other cue. Specifically, we derived a fMRIa pattern distinction score, which was quantified as within-cue similarity (cross-validated Pearson correlation ^25^ between fMRIa vectors of the same cue type) minus between-cue similarity (Pearson correlation between fMRIa vectors of different cue types). A pattern distinction score significantly greater than 0 would indicate that the across-voxel fMRIa pattern was distinct between the two cue types. The analysis was based on objective location, given the previous finding that fMRIa was mainly driven by objective location instead of subjective location.

**Figure 4.**
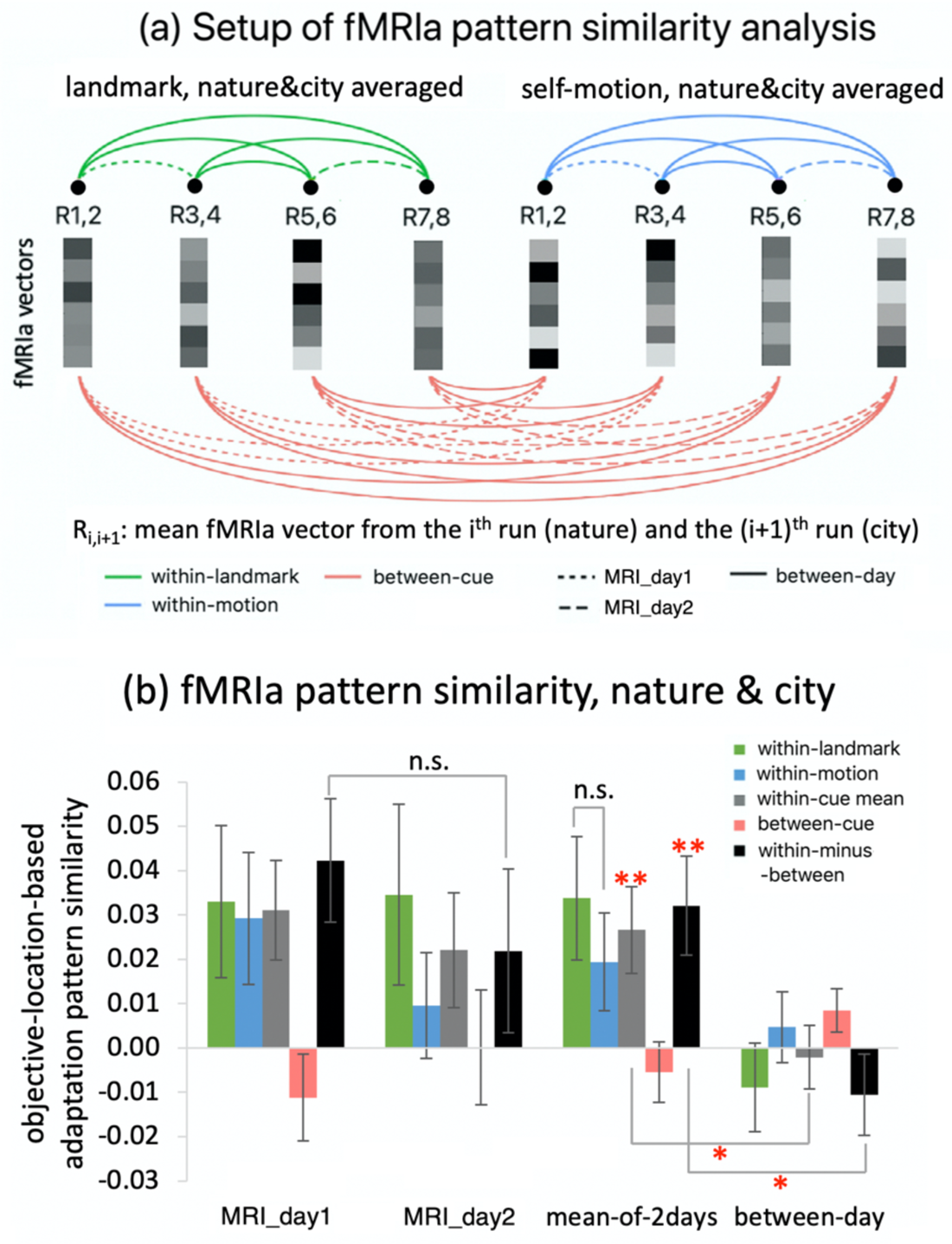
fMRIa pattern similarity analysis in retrosplenial cortex. (a) Setup of the fMRIa pattern similarity analysis. First, runs were ordered chronologically for each cue type. Odd-numbered runs belonged to the nature environment, and even-numbered runs belonged to the city environment. Second, to minimize any subtle influences of environment, for each cue type, the fMRIa vectors estimated from the two adjacent runs from the two different environments were averaged to obtain the mean fMRIa vectors; that is, for each cue type, the fMRIa vectors from an odd-numbered run and the subsequent even-numbered run were averaged (e.g., 1^st^ run and 2^nd^ run were averaged, 3^rd^ run and 4^th^ run were averaged, etc.). In particular, R_i,i+1_ refers to the mean fMRIa vector averaged from the i^th^ run (nature) and the (i+1)^th^ run (city). Next, all the mean fMRIa vectors were paired up to one another, resulting in 3×3 = 9 different types of pairing: cue type (within-landmark vs. within-motion vs. between-cue) × day type (within-day1 vs. within-day2 vs. between-days). (b) Results of the fMRIa pattern similarity analysis. Objective-location-based fMRIa pattern similarity is plotted as a function of cue type and day. See more details of the analysis in STAR Methods. n.s. denotes p_1-tailed/2-tailed_ > 0.1, * denotes p_1-tailed/2-tailed_ < 0.05, and ** denotes p_1-tailed/2-tailed_ < 0.01; + denotes p_1-tailed/2-tailed_ < 0.1.

As shown in Figure 4b, because the within-day pattern distinction score was significantly greater than the between-day distinction score (t(19) = 2.825, p_2-tailed_ = 0.011, BF_10_ = 4.799), we analyzed them separately. The within-day pattern distinction score was significantly greater than 0 (t(19) = 2.885, p_1-tailed_ = 0.005, BF_10_ = 10.625), because the within-cue similarity was significantly positive (t(19) = 2.708, p_1-tailed_ = 0.007, BF_10_ = 7.694) while the between-cue similarity was not (t(19) = −0.807, p_1-tailed_ = 0.785, BF_10_ = 0.141). These results mean that while the fMRIa pattern remained stable for a given cue type, it differed between the two cue types.

In contrast, the between-day pattern distinction score was not significantly greater than 0 (t(19) = −1.145, p_1-tailed_ = 0.867, BF_10_ = 0.120), because neither the within-cue similarity nor the between-cue similarity was significantly greater than 0 (within-cue, t(19) = −0.292, p_1-tailed_ = 0.613, BF_10_ = 0.189; between-cue, t(19) = 1.717, p_1-tailed_ = 0.051, BF_10_ = 1.513). Note that the within-cue similarity was also significantly greater for within-day than between-day (t(19) = 2.368, p_2-tailed_ = 0.029, BF_10_ = 2.162).

Further analyses showed that the same pattern of results existed in each environment, meaning that the above-mentioned results were not solely driven by a single environment (Figure S2a). More control analyses revealed the same pattern of results, when we analyzed the subjective-location-based fMRIa and when the inter-location distance was modeled continuously in both cue conditions (Figure S2b).

To summarize, the fMRIa patterns were correlated within the same cue type but uncorrelated between different cue types. This effect occurred within the same scanning day, and was driven by the temporally stable within-cue fMRIa patterns. Taken together, these results suggest that although RSC encoded spatial distances for both landmarks and self-motion cues in the form of fMRIa, the underlying spatial representations were distinct between the two cue types.

### RSC showed MVPS-based spatial distance coding for both cue types, which was cue-independent and mainly driven by subjective location

Previous results have shown that RSC activation showed adaptation as a function of spatial distance. Here, we investigated whether RSC also contained a similar spatial coding in the form of multi-voxel pattern similarity (MVPS) (Figure 5a). The rationale is that locations closer to each other in space should evoke more similar neural representations as indexed by the multi-voxel activation pattern. First, we calculated MVPS between two test locations as correlational similarity between their corresponding across-voxel activation patterns. Next, a spatial information score was obtained by correlating activation pattern similarity with inter-location distance. A spatial information score greater than 0 would indicate that distances between test locations were encoded in the brain activity. We calculated spatial information scores both within and between the two cue types. The within-cue spatial information scores informed whether distances were encoded for a given cue type. The between-cue spatial information score informed whether the spatial coding was generalizable between cue types, which would be indicative of common spatial representations for both cue types. For all the three measurements, we modeled spatial distances among test locations in a continuous manner by default (STAR Methods, MVPS-GLM1).

**Figure 5.**
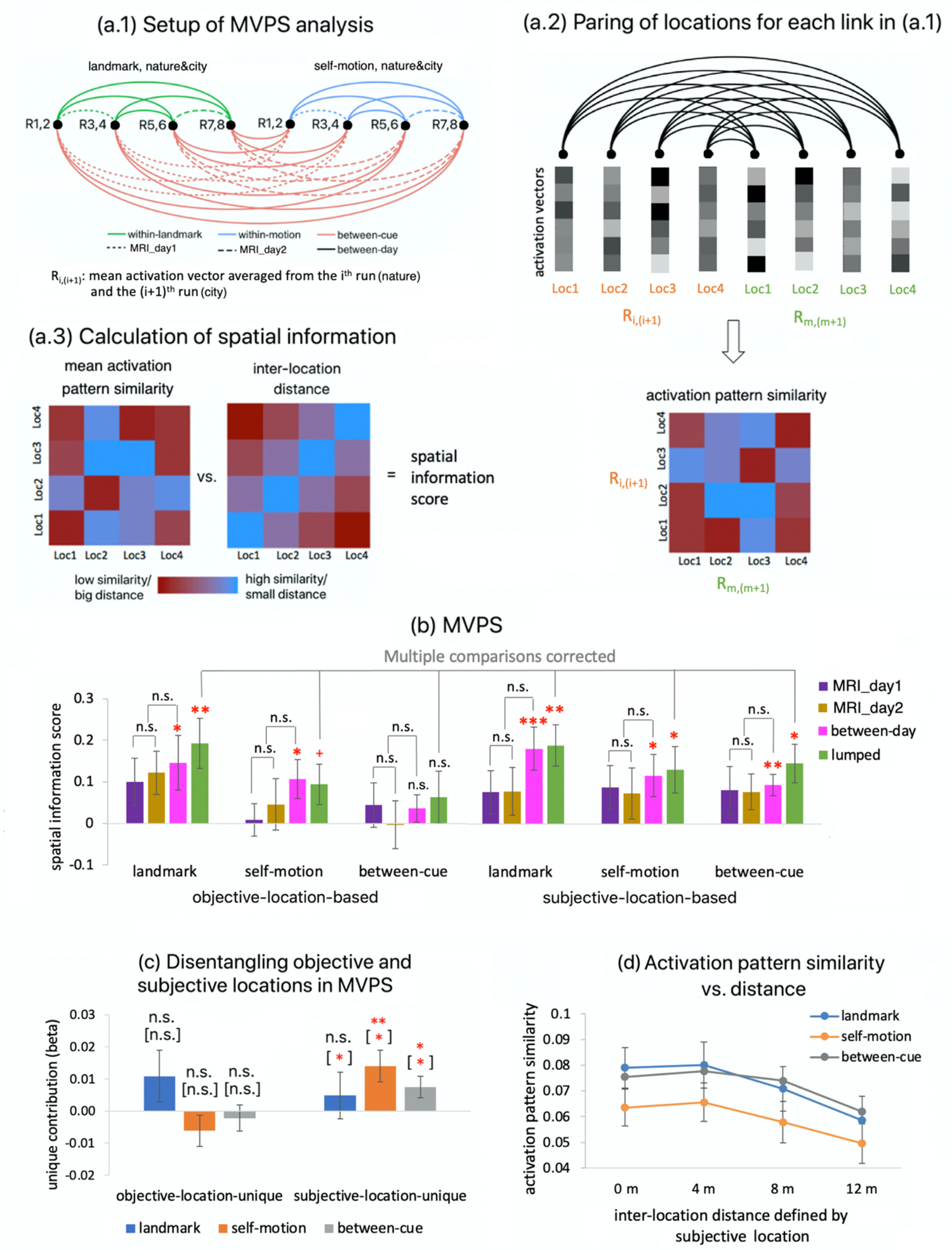
MVPS analysis in retrosplenial cortex. (a) (c) Setup of the MVPS analysis. (a.1) Pairing of the runs. Similar to the fMRIa pattern similarity analysis (Figure 4a), runs were ordered chronologically for each cue type. Odd-numbered runs belonged to the nature environment, and even-numbered runs belonged to the city environment. To minimize any subtle influences of environment, for each cue type, the activation vectors estimated from the two adjacent runs from the two different environments were averaged location by location. (a.2) For each link in (a.1), there are four different mean activation vectors corresponding to the four test locations (i.e., Loc1, Loc2, Loc3, Loc4) in each of the two nodes (i.e., R_i,(i+1)_ and R_m,(m+1)_, in which the subscripts ‘i’ and ‘m’ denote any odd numbers from 1 to 7). In (a.2), calculating the representational similarities between pairwise locations resulted in the 4×4 activation pattern similarity matrix for the link. (a.3) We obtained the 4×4 mean activation pattern similarity matrix by averaging all the similarity matrices across all links in (a.1). Inter-location distances could be defined by either objective or subjective locations. The spatial information score was calculated as the Pearson R correlation between the mean activation pattern similarity matrix and the inter-location distance matrix (Fisher-transformed and reversed in sign). (b) (d) Spatial information score is plotted as a function of cue type (landmark vs. self-motion vs. between-cue), location type (objective vs. subjective location), and day type (MRI_day1 vs. MRI_day2). The lumped spatial information score was calculated regardless of day type (yellow bars). Displayed significance results were corrected for multiple comparisons across the six tests on the lumped spatial information score (yellow bars), using the permutation-based Holm-Bonferroni procedure (STAR Methods). (c) (e) Unique contributions of objective and subjective locations were disentangled in MVPS calculations. Significance levels displayed in the brackets refer to results when participants with behavioral accuracy > 90% were excluded from analysis. (d) (f) To visualize MVPS, activation pattern similarity is plotted as a function of inter-location distance defined by subjective locations for landmarks, self-motion cues, and between cue types. See more details of the analysis in STAR Methods. * denotes p_1-tailed/2_tailed_< 0.05, and ** denotes p_1-tailed/2-tailed_ < 0.01; + denotes p_1-tailed/2-tailed_ < 0.1.

We calculated spatial information scores based on objective and subjective locations and tested them against 0 using one-tailed simple t tests, with the familywise type I error controlled at 0.05 using the permutation-based Holm-Bonferroni procedure for the six individual t tests (measurement (landmark vs. self-motion vs. between-cue) × location type (objective vs. subjective) (STAR Methods). As shown in Figure 5b and Table 1, when objective locations were modeled, spatial information scores were significant in the landmark condition (p_corrected_ = 0.010) but neither in the self-motion condition (p_corrected_ = 0.067) nor between cue types (p_corrected_ = 0.159). When subjective locations were modeled, spatial information scores were significant in both the landmark (p_corrected_ = 0.002) and the self-motion condition (p_corrected_ = 0.046). Critically, the between-cue spatial information score was also significant (p_corrected_ = 0.010). Additional analyses showed that day (Figure 5b) and environment (Table S1) did not affect subjective-location-based MVPS, which was also generalizable between days (Figure 5b) and environments for all three measurements (Table S1). These results suggest that overall MVPS was associated more strongly with subjective than objective location.

To rigorously disentangle objective and subjective location, we directly compared objective-location-defined and subjective-location-defined spatial distances when computing the spatial information scores (STAR Methods, MVPS-GLM2). As shown in Figure 5c and Table 1, the unique contribution of objective location was not significant for all the three measurements (ps_1-tailed_ > 0.09). The unique contribution of subjective location was significant for self-motion cues (p_1-tailed_ = 0.006) and between cue types (p_1-tailed_ = 0.019), but not for the landmarks (p_1-tailed_ = 0.254). When excluding participants with high behavioral accuracies (> 90%) that could cause unreliable estimates, the unique contribution of subjective location was significant for all three measurements (ps_1-tailed_ < 0.03), while the unique contribution of objective location remained non-significant (ps_1-tailed_ > 0.15). These results showed that MVPS was predominantly driven by subjective rather than objective location. These subjective-location-based MVPS effects are visualized in Figure 5d, which shows that activation pattern similarity decreased in a linear manner as inter-location distance increased for landmarks, self-motion cues, and between cue types.

Finally, we conducted the searchlight analysis to investigate MVPS in the entire-volume (Figure S3 & Table S2). In posterior cingulate areas (including RSC proper and the putative retrosplenial complex), MVPS was generally stronger when based on subjective than subjective location in all the three measurements. This is most obvious for self-motion cues and between cue types. This trend also existed in other brain regions, e.g., precuneus, middle occipital gyrus, middle temporal gyrus, and angular gyrus.

To summarize, we found that i) RSC encoded spatial distances for both cue types in the form of MVPS, ii) the coding was mainly driven by subjective rather than objective location, and iii) the coding was generalizable between the cues. Together, these results suggest cue-independent spatial representations in RSC, which also seemed to be cue-invariant, because the spatial information score did not differ among landmarks, self-motion cues, and between cue types (ps > 0.4, BFs_10_ < 0.33).

### Neural space reconstructed from MVPS in RSC resembled navigation behavior

The preceding analyses showed that RSC contained fMRIa-based cue-specific and MVPS-based cue-independent spatial representations. However, a major limitation of both analyses is that different location pairs with the same inter-location distance value were treated equally, which could have obscured potential subtle aspects of the underlying neural representations as suggested by participant’s behavioral performance pattern (Figure 2). Therefore, to better characterize the spatial codes in RSC and their relations to participants’ behavior, we applied a neural space reconstruction analysis and recovered the entire neural space with positional estimates for all the four test locations. The neural space was reconstructed based on neural distances between the four test locations defined by objective location, which was then compared to participants’ behavior and the original physical space (Figure 6a; STAR Methods). Resemblance with the behavioral pattern would indicate that imperfections of the neural representations in RSC for the external physical space might have mediated the behavioral mistakes participants made, whereas resemblance with the original physical space would indicate that the physical space was represented faithfully in RSC.

**Figure 6.**
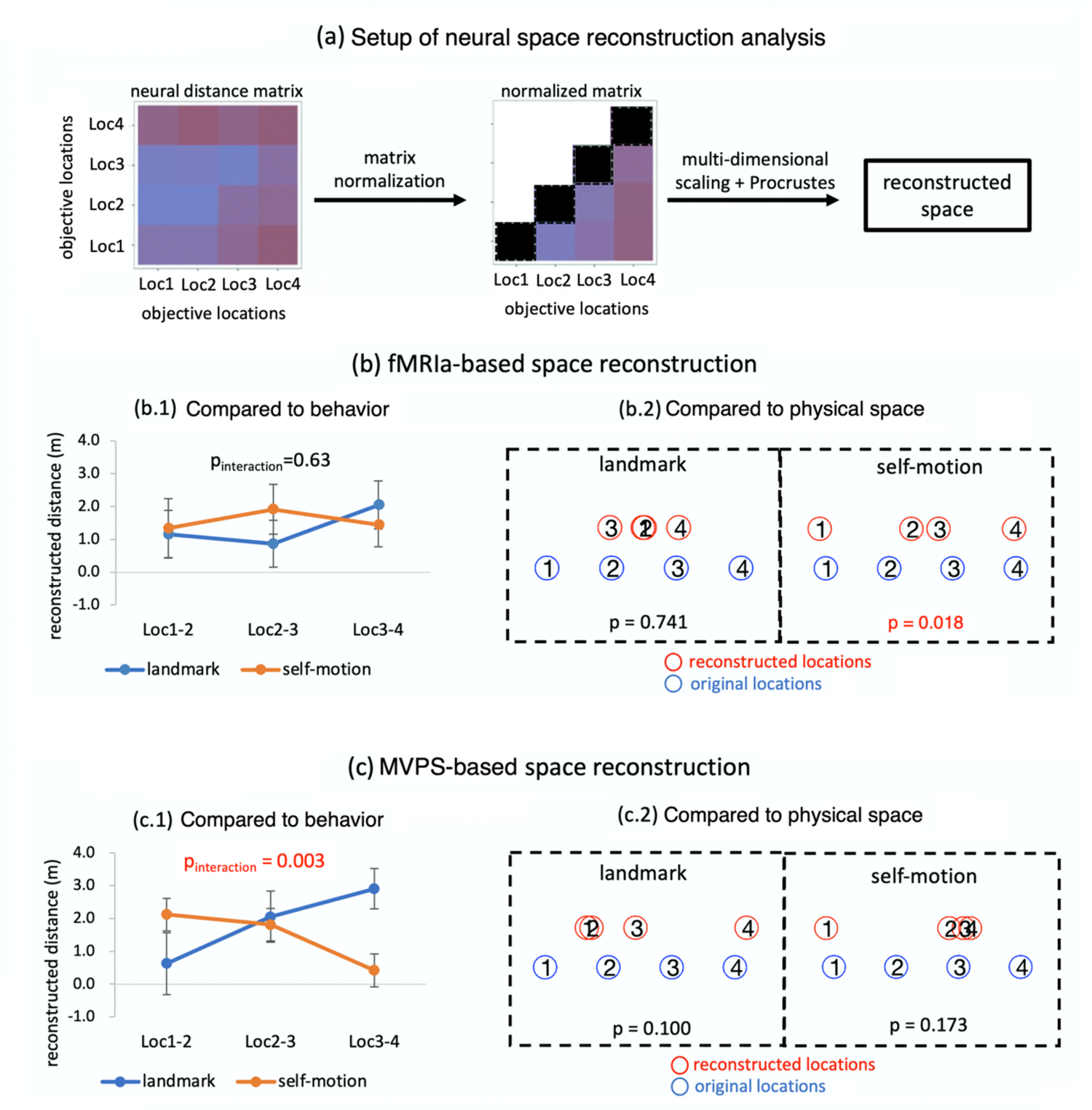
Neural space reconstruction analysis in retrosplenial cortex. (a) Setup of the analysis. For both fMRIa and MVPS, first a 4×4 neural distance matrix was constructed, with the elements denoting pairwise neural distances among the four test locations (defined by objective location). Next, this matrix was normalized, so all the elements were within the range [0, 1], and the four on-diagonal elements were manually set to 0 (dark cells). Third, the normalized neural distance matrix was submitted to the multi-dimensional scaling and then Procrustes analysis to obtain the reconstructed space. See STAR Methods for more details. (b) Results based on fMRIa. (b.1) The pattern of the reconstructed space did not resemble the observed behavioral pattern (Figure 2). The reconstructed distance between adjacent locations is plotted as a function of location pair and cue type, and the interaction between the linear trend of location pair and cue type was not significant. (b.2) Nonparametric permutation tests based on the grand group-level neural distance matrix revealed that the recovered neural space significantly resembled the original physical space for self-motion cues, but not landmarks (STAR Methods). (c) Results based on MVPS. (c.1) The structure of the reconstructed space resembled the observed behavioral pattern (Figure 2). The reconstructed distance between adjacent locations is plotted as a function of location pair and cue type, and the interaction between the linear trend of location pair and cue type was significant. (c.2) Nonparametric permutation tests based on the grand neural distance matrix revealed that the recovered neural space did not significantly resemble the original physical space for either cue type. See more details of the analysis in STAR Methods.

For fMRIa, we first submitted the participant-specific reconstructed neural distances between adjacent locations to a repeated-measures ANOVA, with cue type (landmark vs. self-motion) and adjacent location pair (Loc1-2 vs. Loc2-3 vs. Loc3-4) as independent variables (Figure 6b.1). Unlike the behavioral pattern, the interaction effect between the linear trend of location pair and cue type was not significant (F(1,19) = 0.239, p_1-tailed_ = 0.630, η_p_^2^ = 0.012). Next, to assess the similarity with the original physical space, we conducted a permutation-based test on the group-level neural distance matrix (Figure 6b.2). In the landmark condition, the neural space did not significantly resemble the original space (p = 0.741), with some locations even swapped in order (e.g., Loc3 was to the left of Loc1). This echoes with the earlier observation that in the landmark condition, the repetition suppression effect seemed to only discriminate between same and different locations, but not between different non-zero inter-location distances (Figure 3c). On the contrary, in the self-motion condition, the neural space significantly resembled the original physical space (p_1-tailed_ = 0.018). This also echoes with the earlier observation that in the self-motion condition, the repetition suppression effect appeared to occur in a linear manner over the entire range of inter-location distance (Figure 3c). In brief, fMRIa-based neural space (i) bore no similarities to behavior in the two cue conditions and (ii) significantly resembled the original physical space in the self-motion condition.

For MVPS, first, we analyzed participant-specific neural distances, and observed significant interaction between cue type and the linear trend of location pair (F(1,19) = 12.016, p = 0.003, η_p_^2^ = 0.387): the neural distance between adjacent locations decreased as the locations became farther away from the landmark in the landmark condition, whereas the pattern was reversed in the self-motion condition (Figure 6c.1). This is parallel to the behavioral performance pattern (Figure 2). Next, we analyzed the group-level neural distance matrix, and found that the recovered neural space did not significantly resemble the original physical space in either cue condition (landmark, p_1-tailed_ = 0.100; self-motion, p_1-tailed_ = 0.173; Figure 6c.2); furthermore, the group-level neural space (Figure 6c.2) exhibited a structure qualitatively similar to the behavioral performance pattern (Figure 2).

To summarize, the fMRIa-based neural spaces did not resemble participants’ behavior. Furthermore, in the self-motion condition, the fMRIa-based neural space resembled the original physical space, suggesting a map-like spatial code that maintained Euclidean distances among test locations without salient spatial distortions. In contrast, the MVPS-based neural spaces exhibited a pattern similar to participants’ behavior and did not resemble the original physical space. Taken together, compared to fMRIa, MVPS was more closely associated with behavior along the stimulus-response spectrum, which is consistent with the preceding observation that MVPS was fitted better by subjective-location-defined than objective-location-defined spatial distances.

### Hippocampus contained a spatial coding scheme similar to RSC

We found that the hippocampus, whose trial-by-trial activation was strongly correlated with RSC (Figure S4), also showed a spatial coding scheme similar to that of RSC. In particular, the hippocampus exhibited a trend towards fMRIa-based cue-specific spatial representations (Figure S5), and MVPS-based cue-independent spatial representations that resembled participants’ behavior (Figure S6). However, compared to RSC, these effects in the hippocampus were evidently reduced in magnitudes, and objective location and subjective location were less well dissociated in the neural coding (e.g., Figure S6b&e).

## DISCUSSION

The current study investigated whether landmark-based navigation and path integration recruit cue-specific or cue-independent spatial representations in the human RSC and hippocampus. Participants completed a spatial navigation task on a linear track, in which the use of landmarks and self-motion cues was dissociated, but they used these cues to encode and retrieve the same set of spatial locations. In RSC, we found clear evidence for the existence of both cue-specific and cue-independent spatial representations. Cue-specific spatial representations were revealed through fMRIa: while RSC displayed repetition suppression for both landmarks and self-motion cues, the distributed fMRIa patterns were distinct between cue types. Cue-independent spatial representations were revealed through MVPS, in that the similarity of multi-voxel activation patterns between two locations – defined by the same or different cue types – decreased as the inter-location distance increased. Additionally, while fMRIa-based spatial representations were more related to objective sensory inputs, MVPS-based spatial representations were more strongly associated with behavior that differed from the sensory inputs. The hippocampus exhibited strong functional connectivity with RSC and showed a similar spatial coding scheme, but the effects were generally weaker. To our knowledge, the current study is the first demonstration in humans that both types of spatial representations co-existed in the same brain region while participants were performing a navigation task in the same spatial context.

One prominent feature of the current study is that landmarks and self-motion cues were clearly dissociated, which is evident in the differential behavioral profiles of the two cue conditions. Specifically, behavioral performance increased as the test location got closer to the landmark in the landmark condition, whereas the opposite pattern was observed in the self-motion condition. This is because while the spatial precision afforded by the landmark (i.e., the anchoring point of landmark-based navigation) deteriorates as the location becomes farther away from it ^3,26^, path integration gets noisier as the navigator travelled along the path and away from its anchoring point – the fixed starting position ^27^. This finding is broadly consistent with previous studies showing a relative independence of path integration and landmark-based navigation in behavior ^1, 2^, which suggests that our cue dissociation manipulation successfully elicited distinct navigational strategies in the two different cue conditions. On the contrary, spatial cues were not clearly dissociated in most of the previous related studies ^10–12, 14, 16^. For example, in Huffman and Ekstrom’s human fMRI study ^16^, visual information was present in all conditions that differed in the degree of body-based self-motion cues, raising the possibility that the reported cue-independent neural representations may have been driven by the ever-present visual information.

The simultaneous investigation of fMRIa and MVPS was the key factor to reveal both cue-specific and cue-independent spatial representations in RSC (and potentially in the hippocampus as well). When the sensory inputs for encoding the same physical space changed, previous studies have observed either cue-specific spatial representations in the hippocampus ^9^, or cue-independent spatial representations in RSC ^14^ and in the brain-wide functional connectivity pattern ^16^. A recent study found that whether altering spatial inputs invoked cue-specific or cue-independent spatial representations in the hippocampus depended on whether different cue types were congruent in defining the reward location ^13^. Taken together, previous studies have observed either cue-specific or cue-independent spatial representations but not both at the same time in the same spatial context. Note that in the current study, participants always performed the same navigation task by encoding and retrieving the same spatial locations in the same environment with the same reward configuration; the sole difference between different cue conditions was the type of spatial information available. Therefore, our observation of concurrent cue-specific and cue-independent spatial representations could not have been confounded by factors like task requirement or reward setup. Moreover, our findings suggest that previous studies reporting cue-independent spatial representations might have missed parallel cue-specific representations reflected in a different form of neural activity ^13, 14, 16^. For this reason, the current study highlights the importance of investigating complementary neural phenomena to obtain a more complete understanding of the neural representations underlying cognitive maps.

Cue-specific and cue-independent representations were revealed by fMRIa and MVPS, respectively. These two approaches can yield inconsistent results ^17, 28–32^, which may indicate that they interrogate different aspects of neural operations. One hypothesis posits that fMRIa is related to the processing of neuronal inputs, whereas MVPS reflects neuronal output ^17^. Consistent with this hypothesis, neuronal adaptation (i.e., reduction in neural responses to the same or a similar stimulus) in the macaque inferior temporal cortex was smaller between two different stimuli – compared to two identical stimuli –, even though both stimuli activated the neuron to the same extent ^33–35^. This stimulus dependency indicates that neuronal adaptation may occur locally at the level of the synapses onto the neuron ^35^. Consistently, fMRIa seems to be relatively independent of top-down cognitive operations such as task requirement ^36^ and attentional state ^33^ that typically affect behavior. In contrast, previous human neuroimaging studies frequently observed tight relationships between MVPS and overt behavior ^29^, e.g., more distinct MVPS-based neural representations of different items correspond to better discrimination performance ^37–40^. Consistently, the collective activity of the place cell population in the rodent hippocampus encodes the animal’s subjective recognition of the reward location, regardless of whether it matched the true reward location or not ^41^. This indicates that the collective neuronal output of hippocampal place cells is rather linked to response output instead of stimulus input.

Our results also provide support for the input-versus-output hypothesis, in that fMRIa and MVPS effects in RSC were associated with objective and subjective locations, respectively. What does this reveal about the underlying neural mechanisms in RSC? First, fMRIa patterns based on objective locations were spatially dissociated between landmarks and self-motion cues, which would be consistent with separate location-sensitive neuronal subpopulations that were driven by sensory inputs from the two cue types, respectively. These subpopulations should display adaptation to the stimulation from external spatial inputs. Second, MVPS based on subjective locations was spatially generalizable between cue types, which would be consistent with a location-sensitive neuronal subpopulation whose ensemble activity represented the navigator’s subjective location in a cue-independent manner. Importantly, this particular subpopulation should not display adaptation, because the fMRIa patterns would otherwise have shown spatial overlap between the cue types and hence eliminated the cue-specificity we observed in the distributed fMRIa patterns. Finally, additional analyses showed that MVPS and fMRIa were relatively independent at the voxel level (Figure S7), indicating that the non-adapting subpopulation representing subjective locations was probably anatomically separable from the adapting subpopulations encoding objective locations.

This interpretation is corroborated by recent observations in rodents. Brennan et al. (2019) discovered different types of cells in the rodent RSC, with one cell type adapting to external stimulation and the other type showing no adaptation but firing persistently in the presence of continued stimulation ^42^. Importantly, their modeling work suggests that it is the activity of the non-adapting cells – but not the adapting cells – that represents the animal’s current head direction when the animal remains still, a scenario similar to the navigation task used in the current study (i.e., we analyzed fMRI data acquired from when the participants’ first-person perspective was fixed at the test locations for 4 seconds). Furthermore, Fischer et al. (2019) found in rodents that V1 projections to RSC displayed similar spatial tuning as the place-cell-like cells in RSC, but with less modulation of the animal’s navigation state (i.e., active vs. passive navigation) ^43^. This echoes with our observation that cue-specific fMRIa effects were tied closely to objective locations and less so to overt navigation behaviors. Taken together, findings of these rodent studies accord with our interpretation that different neuronal subpopulations may serve different computational purposes in RSC.

The functional properties of RSC put it in a good position to support participants’ navigation behavior in the current study. The completion of the location identification task recruited two main cognitive components: a long-term memory component, which corresponds to the cue-independent memory traces of the four test locations learned over time, and a perception component, which corresponds to perceiving instantaneous cue-specific sensory inputs. Participants had to compare the two components to judge which of the four stored memory traces best matched the current sensory inputs. RSC subserves long-term spatial memories ^44–46^ and receives projections from brain regions that process a variety of sensory information, including several visual areas (incl. V1, V3 and V4 ^47^), thalamus ^48^, and areas in the medial temporal lobe ^49^. Therefore, RSC appears to mediate the interaction between long-term memory (likely reflected in MVPS) and perception (likely reflected in fMRIa) to facilitate cognitive map formation ^5–7^, e.g., by integrating different spatial inputs with preexisting memory traces to construct coherent spatial representations. This might explain why RSC’s neuronal output was closely related to the response output (i.e., participants’ behavior), and why RSC’s spatial coding scheme corresponded to the input-versus-output hypothesis in the current study.

Finally, we found that RSC and the hippocampus showed strong functional connectivity along with similar spatial coding schemes, which accords with recent findings in rodents ^14, 15, 50, 51^. Note that like RSC, the hippocampus is also well-suited to mediate the interaction between long-term memory and perception, because it is crucial for memory formation ^52, 53^ and also receives multisensory inputs via the entorhinal cortex ^54^. However, although BOLD signal quality in terms of the temporal signal-to-noise ratio was comparable between the two regions (Table S4), effects were generally weaker in the hippocampus. This difference could be related to the memory stage our participants were at during the scanning. Past work has indicated that hippocampal and RSC activity reflects the learning rate and the learning amount, respectively, so that the hippocampal involvement decreases whereas RSC involvement increases as spatial memories are being formed ^44, 55^. In addition, spatial representations appeared to shift from the hippocampus to RSC during the course of memory formation ^45^. Our results showed that compared to the behavioral training day prior to the MRI scanning, participants’ performance improved on the first scanning day but remained unchanged between the two scanning days (Figure 2 & Table S5), indicating that participants might have already reached late stages of memory formation during scanning. Consistently, additional fMRI analyses revealed that RSC, but not the hippocampus, was more activated during successful than failed trials (Table S6). This might also explain why we barely observed fMRIa in the entorhinal cortex, except that in the posterior-medial entorhinal cortex the fMRIa-based neural space resembled the original physical space (Table S3). This finding is inconsistent with our previous report of fMRIa-based distance coding in the entorhinal cortex for both landmarks and self-motion cues ^3^. Considering that the hippocampus receives sensory information from cortical areas via the entorhinal cortex ^54^, it is conceivable that the entorhinal cortex should be minimally recruited if the downstream area hippocampus was not much involved in the task. Consistent with the interpretation, we found that along with the hippocampus, the entorhinal subregions contributed to successful navigation in our previous study but not in the current study (Table S6). Therefore, future work is needed to investigate the temporal dynamics between the hippocampal formation and RSC in representing spatial information at different memory stages.

## CONCLUSION

In this study, we investigated a core question in spatial navigation– whether landmark-based navigation and path integration recruit common or distinct spatial representations in the brain. We demonstrated the coexistence of cue-specific and cue-independent spatial representations in the human RSC. Furthermore, by establishing a human fMRI paradigm highly similar to paradigms widely used in non-human animal studies, we hope the current study will facilitate inter-species comparisons in spatial navigation.

## STAR Methods

### EXPERIMENTAL SETUP AND SUBJECT DETAILS

#### Participants

Twenty healthy adult volunteers from the Magdeburg community participated in this experiment (10 male; mean age = 25.35 year old, standard deviation of age = 3.91 year). All participants were right-handed, had normal or corrected-to-normal vision, and had no neurological diseases. Three additional participants were tested but were excluded from data analysis, either because they dropped out in the middle of the experiment or because the fMRI data were corrupted by technical problems. All participants gave informed consent prior to the experiment and received monetary compensation after the experiment. The experiment was approved by the Ethics Committee of the University of Magdeburg.

#### Stimuli and navigation task

Virtual environments were created and rendered in Worldviz 5.0 (https://www.worldviz.com). There were two different virtual environments, a city environment and a nature environment (Figure 1a). These two environments had different background views and different ground textures. A linear track was included in both environments. The linear tracks were covered with the same texture but rendered in different colors in the two environments. The linear tracks shared the same object configuration in the two environments (Figure 1b). Three arrows and a tree were positioned at the object layout on the track. The tree was slightly to the left from the imagery midline of the linear track (= 0.5 m). In between the arrows and the tree were four balls of different colors positioned at four test locations. The four test locations were evenly spaced in the linear track with intervals of 4 m. To further distinguish the two environments, the order of the four balls was reversed between the two environments, but they occupied the same four test locations in both environments. Both the arrows and the tree were identical but rendered in different colors in different environments.

#### Learning task

Participants used a MRI-compatible joystick to navigate around in the virtual environments and give responses. Participants were trained to learn four test locations that were evenly spaced on the linear track (Figure 1a). Four balls of different color were positioned at the four test locations. Participants needed to remember the colors of the balls associated with the test locations (see the video – the part “LEARNING”).

#### Test: Location identification task

In the ‘location identification task’, the participant was passively transported to one of the four test locations, and was required to recall the color of the ball positioned at this test location, while the ball remained invisible throughout the trial. The time course of a trial is depicted in Figure 1b (also see the video – the part “TEST: location identification task”). In each trial, the starting position of the passive movement was randomly sampled from a uniform distribution U(−18m, −4m) on a trial-by-trial basis (Figure 1a). Once the passive movement had stopped, the participant’s first-person perspective was fixed at the test location for 4s, after which they had to report the color of the ball positioned at the location they thought they were now occupying. Importantly, the order of the four options appearing on the screen was randomized from trial to trial, and a randomly selected option was highlighted as the initial answer before the participant started to make response. In addition, participants pressed only one particular button on the joystick to switch among the options in a loop. In this way, each test location was not associated with any fixed option position on the screen or with any consistent pattern of finger movement on the joystick. To prevent pure timing or counting strategies, the movement speed was randomly sampled from a uniform distribution U(2 m/s, 5 m/s) on a trial-by-trial basis. Accuracy was emphasized, but participants were instructed to not spend longer time than necessary.

The use of self-motion cues and landmark cues was dissociated in the task, in a way similar to Chen et al. (2019) with minor adjustments ^3^. This manipulation followed the logic of dissociation of landmark and self-motion cues in established behavioral paradigms ^1, 56, 57^. In the self-motion condition, the arrows and the linear track texture were both visible. Because the arrows could serve as the anchoring point for path integration on travelled distance, the participant could perform path integration on travelled distance based on optic flow after he/she had passed the arrows. The landmark was not visible, meaning that landmark-based navigation was eliminated. To prevent participants from associating the test locations with any spatially isolated features on the ground, which would resemble the landmark-based navigation strategy (e.g., the red ball’s position was always within the brightest patch of the ground), both the texture of the linear track and the texture of the floor outside of the linear track were randomly shifted in position along the long dimension of the track from trial to trial based on a uniform distribution U(−50m, 50m).

On the contrary, in the landmark condition, the landmark was visible, meaning that the participant could rely on the landmark for localization. To eliminate path integration, the arrows were invisible, and the ground of the linear track remained blank to remove the texture information. Although there was still peripheral optical flow stemmed from the floor texture outside of the linear track, since the starting position of the passive movement was randomized on a trial-by-trial basis and the anchoring point for path integration (i.e., the arrows) was invisible, the participant could not perform path integration to solve the task, i.e., the participant would not know how far he/she needed to travel to reach a ball location. The cue manipulation in the landmark condition is analogous to the disorientation manipulation typically used to eliminate self-motion information in spatial navigation studies ^58, 59^.

### Experimental procedure

The experiment took place on three consecutive days, with behavioral training on the 1^st^ day (Pre-scan_day) and MRI scanning on the 2^nd^ day (MRI_day1) and 3^rd^ day (MRI_day2) (Figure 1c). The time interval between Pre-scan_day and MRI_day1 varied between 1-17 days (mean=2.75), and the time interval between MRI_day1 and MRI_day2 varied between 1-17 days (mean = 4.35). For two participants, the time interval between the two scanning days was 17 days, due to the restricted availability of the participants and the MRI scanner.

#### Behavioral training (Pre-scan_day)

The behavioral training allowed the participants to get familiar with the virtual reality environment and to learn the four test locations. The training consisted of three parts. Each part had a learning stage and a test stage. During the learning stage (see the video – the part “LEARNING”), first, participants learned the colors of the balls positioned at the four test locations and were tested on their memory of the colors. Next, they learned the locations of the four balls. In each trial, one ball was displayed, and the participant actively moved from a randomized starting position to the ball’s location. Both the landmark and self-motion cues were available, meaning the arrows, the tree, and the ground texture of the linear track were all visible. Each ball was learned twice, with the order of the four balls counterbalanced. The learning stage was performed twice for each environment, with the order of the two environments counterbalanced. The learning stage was identical for all the three parts in the Pre-scan day. During the test stage, the participant was tested in the ‘location identification task’, as described in the preceding section (Figure 1b; also see the video – the part “TEST: location identification task”). There were four blocks in total (counterbalanced), corresponding to the four combinations of environment (city vs. nature) and cue condition (self-motion vs. landmark). In the first part, each block had 4 trials, corresponding to the four ball locations (counterbalanced). In the second and the third parts, during the test stage, each block had 16 trials, with 4 trials for each ball location (counterbalanced). The experimenter carefully instructed the participants from the beginning to the end during the first part of the Pre-scan training. For the remaining two parts of the training, participants were left alone to perform the tasks, but were attended by the experimenter when needed.

#### MRI scanning (MRI_day1 & MRI_day2)

The two scanning day sessions shared the same procedure. On each scanning day, we first re-familiarized participants with the task by requiring them to practice the task while they were undergoing structural scanning inside the scanner. The practice stage was exactly the same as the first part in the behavioral training day (Pre-scan_day). This practice stage lasted about 5 minutes and was not analyzed further. During the subsequent functional scanning, participants performed the ‘location identification task’ (Figure 1b; also see the video – the part “TEST: location identification task”). On each scanning day, there were eight runs in total, with two runs for each of the four combinations of environment (city vs. nature) and cue condition (self-motion vs. landmark). The eight runs were organized in two blocks, and in each block, each of the four runs corresponded to one of the four condition combinations. In each block, the four condition combinations were semi-randomized in order, using Latin square designs and with the restriction that the combinations occurring in two successive runs must be different within the same day.

We adopted a continuous carry-over design ^18^. We used the eight de Bruijn sequences from our previous study with relatively high detection power and low correlation coefficient^3^. These de Bruijn sequences were generated with 2nd order counterbalancing, using the ‘path-guided’ approach ^19^. In these de Bruijn sequences, the ‘carry-over’ effects (i.e., the influence of a prior item on the brain response to the current item) were counterbalanced, allowing us to investigate fMRI adaptation and multi-voxel pattern similarity simultaneously with the same set of trials ^18, 22^. There were five types of events in each sequence – fixation periods at the four test locations, in which participants stayed at the test locations for 4s, and null events, in which participants fixated their eyes at a cross displayed in the middle of the blank screen. Each de Bruijn sequence contained 25 events in total, with five repetitions for each event type. To allow the hemodynamic response to reach a steady state before the sequence started, we duplicated the very last event in the sequence and placed it at the very beginning. This duplicated event was modeled in the first-level GLMs, but was not included for the analyses of the fMRIa or MVPS effects. Therefore, in each run, there were 20 effective trials in total for the fMRIa and MVPS analyses, with five trials for each of the four test locations. These eight de Bruijn sequences were then randomly assigned to the eight runs in each scanning day for each participant. On each day, the functional MRI scanning lasted up to about 1 hour, and the total scanning time lasted up to about 1.75 hour.

### MRI acquisition

Structural and functional images were acquired in a 7T MR scanner (Siemens, Erlangen, Germany) at the Leibniz Institute for Neurobiology in Magdeburg with a 32-channel head coil (Nova Medical, Wilmington, MA). A high-resolution whole-brain T1-weighted structural scan was acquired with the following MP-RAGE sequence: TR = 1700 ms; TE = 2.01 ms; flip angle = 5°; slices = 176; orientation = sagittal; resolution = 1 mm isotropic. A partial-volume turbo spin echo high-resolution T2-weighted structural scan was acquired perpendicular to the long axis of the hippocampus (TR = 8000 ms; TE = 76 ms; flip angle = 60°; slices = 55; slice thickness = 1 mm; distance factor = 10%; in-plane resolution = 0.4 × 0.4 mm; echo spacing = 15.1 ms, turbo factor = 9, echo trains per slice = 57). Functional scans were acquired with a T2*-weighted 2D echo planar image slab centered on the hippocampus and parallel to its long axis (TR = 2000 ms, TE = 22 ms; flip angle = 85 °; slices = 35; resolution = 1 mm isotropic, parallel imaging with grappa factor 1, echo spacing = 0.82 ms). We also obtained 10 volumes of whole brain functional scans for the purpose of co-registering anatomical masks obtained on the T2-weighted structural scan to functional scans with a MPRAGE sequence (TR = 5000 ms, TE = 22 ms; flip angle = 85 °; slices = 100; resolution = 1.6 mm isotropic). The T1-weighted structural image was bias-corrected in SPM12. Functional scans were motion and distortion corrected online via point spread function mapping ^60^. Functional scans were left spatially unsmoothed. Figure 1d shows the T2-weighted structural scan and a functional scan overlaid on the T1-weighted structural scan for an exemplary participant.

### Anatomical masks for regions of interest

As our regions of interest (ROI), we focused the retrosplenial cortex (RSC) and brain regions in the medial temporal lobe (MTL), including hippocampus, parahippocampal cortex (PHC), entorhinal cortex (EC), and perirhinal cortex (PRC). All the anatomical masks were obtained in the native space of each participant’s structural scans. To illustrate, Figure 1d, Figure S4a, and Figure S5c displays the anatomical masks for an exemplary participant.

The procedure for obtaining the anatomical mask for RSC was identical to that used in a previous study in our lab (Shine et al., 2016). RSC mask was automatically extracted from each participant’ T1-weighted structural scan (bias-corrected in Advanced Normalization Tools (ANTs)) in Freesurfer ^62^, using the ‘recon-all’ command. RSC was defined as the posterior-ventral portion of the cingulate gyrus, which mainly consists of BA29/30. Note that the definition of RSC is anatomically different from the retrosplenial complex, which is a functionally defined region typically extending into the parieto-occipital sulcus ^63^. Although we did not investigate the retrosplenial complex in the ROI-based analyses, we conducted corresponding fMRI analyses to explore in the entire volume that likely included the putative retrosplenial complex (Figure 1d).

Brain regions in MTL were manually segmented in each participant’s T2-weighted structural scan in ITK-SNAP (Yushkevich et al., 2006; http://www.itksnap.org/pmwiki/pmwiki.php), following the protocol developed by Berron, Vieweg and colleagues ^65^. As shown in Figure S5c, the hippocampus was further segmented into different subfields (CA1, CA2, CA3, subiculum (SUB), dentate gyrus (DG), and tail), using the same protocol ^65^. As shown in Figure S4a, EC was further divided into the anterior-lateral subregion (alEC) and the posterior-medial subregion (pmEC), following the procedure developed in our previous study ^3^.

The anatomical mask for RSC was first co-registered to the mean functional scan along with the T1-weighted structural scan in SPM12; then the co-registered anatomical mask was resliced using the nearest-neighbor interpolation, with the mean functional scan as the reference image. The anatomical masks for the MTL regions were co-registered to the mean functional scan of the first scanning day in SPM12, using the same procedure adopted in our previous study ^3^: first, the mean whole-volume functional scan was co-registered to the mean functional scan; second, the T2-weighted structural scan, along with the anatomical masks, were co-registered to the mean whole-volume functional scan obtained from the first step; third, the co-registered anatomical masks were re-sliced using nearest-neighbor interpolation, with the mean functional scan as the reference image.

### STATISTICAL ANALYSIS

#### Behavioral data analyses

We calculated behavioral accuracy based on whether the answer was correct (coded as 1) or not (coded as 0), with a chance level of 0.25. For the two scanning days, the first trial of the sequence in each block was not included in the analysis, because it was not included in the main fMRI analyses and did not appear to differ from other trials in the sequence. In the main text, we focused on behavioral data from the two scanning days (Figure 2). We reported results of the behavioral data from all the three days in the supplemental information (Table S5).

#### Cognitive modeling to recover representational precision from behavior

To dissociate representational precision from response bias in behavioral performance, we applied an extension of signal detection theory to our location identification task with four choices. In the modeling, we included eight free parameters to model i) the four standard deviations of the underlying representations of the four test locations (*S_1_, S_2_, S_3_, S_4_*), ii) the three response criterions (*C_12_, C_23_, C_34_*), and iii) the lapse rate (*lr*). The lapse rate represents the proportion of trials in which participants completely failed in attention and simply chose a response randomly. The centers of the representation distributions (i.e., *μ_1_, μ_2_, μ_3_, μ_4_*) were assumed to be at the true positions of the test locations (i.e., *μ_1_* = −6m, *μ_2_ =* −2m, *μ_3_ =* 2m, and *μ_4_ =* 6m).

In each simulation, we constructed the behavioral confusion matrix, given a set of algorithm-generated values for the eight free parameters. Specifically, for the (1 − lr) proportion of the trials, we randomly sampled a sensory input (*x*) from the normal distribution of the underlying representation corresponding to the test location presented in that trial, *N(μ_r_, S_r_)*. Then, a response R(*x*) was made by comparing the sensory input to the three response criterions:

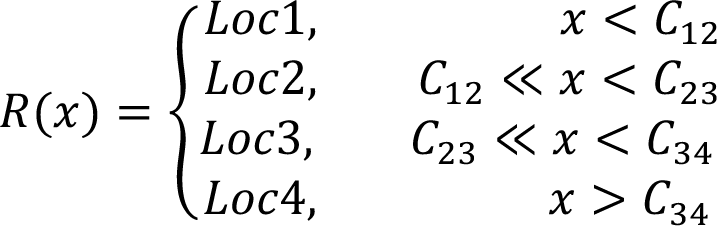

For the remaining *lr* proportion of the trials, we randomly selected one of the four choices as the response, regardless of the current sensory input.

In each simulation, we simulated 10000 trials for each of the four test locations to construct the 4×4 theoretical behavioral confusion matrix. We normalized the theoretical behavioral confusion matrix so that elements in the matrix ranged from 0 to 1, each representing the probability of a response falling to a certain cell of the matrix (i.e., *P_r,c_* – probability of location r recognized as location *c*). We then compared the actual behavioral confusion matrix (Figure 2b) to the theoretical behavioral confusion matrix, by computing the probability of observing each actual response given the theoretical matrix (log-transformed). Finally, we summed the probabilities of all actual responses,

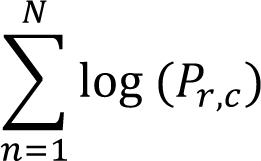

in which n represents the trial number and N represents the total number of trials in the actual experiment. We repeated the simulation to maximize this summed probability (i.e., maximum likelihood estimation). We used the Hooke & Jeeves hill-climbing algorithm for model optimization ^66^, as implemented in Matlab_R2020a. To avoid the potential local-minima problem, the model-fitting procedure was repeated 20 times with randomized starting values for the parameters each time, and the parameter estimates with the best fit were selected (Figure 2c.1).

We performed bootstrapping to estimate variabilities of the estimates for these free parameters. In each iteration, we randomly sampled the same number of responses from the actual responses with replacement for each test location. We then submitted the sampled data to the abovementioned model fitting procedure, and obtained the estimates for the free parameters. The procedure was repeated 600 times, resulting in distributions for all the eight free parameters. 95% confidence intervals of these estimates were obtained from these bootstrapped distributions (i.e., error bars in Figure 2c.2 and Figure 2c.3).

To evaluate how well the model fitted the data (Figure 2c.4), we simulated the behavioral confusion matrix, using the best-fitting values of the eight parameters. We simulated 1000 trials for each test location. We then calculated Pearson correlation between the simulated confusion matrix with the actual confusion matrix. R-squared was taken as a measurement of goodness-of-fit of the model, i.e., the proportion of variance in the data explained by the model. Because the numbers of correct trials and incorrect trials differed dramatically, we evaluated the model fit separately for correct and incorrect trials, as well as separately for the landmark condition and the self-motion condition.

### Functional MRI analyses

#### Univariate analysis of fMRI adaptation

We constructed a first-level general linear model (fMRIa-GLM1) to assess fMRI adaptation (fMRIa). In the model setup, for the regressors that modeled the location occupation periods (Figure 1b, phase 4 ‘location occupation’), we included parametric regressors modeling the modulatory effects of the spatial distance between two successively visited locations. In the self-motion condition, these parametric regressors modeled inter-location distance in a continuous manner by default, i.e., containing values of 0m, 4m, 8m, and 12m. In the landmark condition, these parametric regressors modeled same locations vs. different locations, i.e., containing values of 0 (the two locations were the same) and 1 (the two locations were different), based on a previous report ^22^. The location occupation periods that could not be modeled for fMRIa (i.e., test locations preceded by the null event and the first location occupation event) were modeled with separate regressors. The passive movement phase was modeled with separate regressors, separately for each run and each cue type, but irrespective of the test location. The 16 runs were modeled with separate regressors. The events were convolved with the canonical hemodynamic response function, with the time derivative modeled. Head motion parameters (three rotations and three translations) were entered into the model as nuisance regressors, separately for the 16 runs. Each run was modeled with a constant variable.

We conducted univariate fMRIa analyses based on both objective location (where the participant was actually located) and subjective location (the participant’s response, i.e., where the participant thought he/she was located). Because in the location identification task, the participant was required to explicitly judge the identity of each ball, we could construct the parametric modulation regressors of inter-location distance in terms of subjective location in addition to objective location. For example, if the participant visited Loc1 and Loc3 in two successive trials, but reported “Loc2” and “loc3” in these two trials, the objective-location-based spatial distance was calculated as the physical distance between Loc1 and Loc3 (= 8m), and the subjective-location-based spatial distance was calculated as the physical distance between Loc2 and Loc3 (= 4m). First, we assessed the overall contributions of objective location and subjective location to fMRIa via two versions of fMRIa-GLM1. In fMRIa-GLM1a the parametric regressors of spatial relations were defined by objective location, whereas in fMRIa-GLM1b defined by subjective location. The beta estimates of the parametric regressors represented the overall contributions of objective location in fMRIa-GLM1a and subjective location in fMRIa-GLM1b to fMRIa. Images of the beta estimates for the regressors were left spatially unsmoothed.

At the group-level, in the ROI-based analysis, beta estimates for fMRIa of all the voxels in the ROI were averaged. Then participant-specific beta estimates of the four fMRIa measurements (i.e., location type (objective-location-based vs. subjective-location-based) X cue type (landmark vs. self-motion)) were tested using directional one-sample t tests separately to obtain the uncorrected significance levels (i.e., p_uncorrected_). Next, the four measurements were submitted to a multiple comparisons correction approach that combines the nonparametric permutation-based maximum-t-statistic method ^67^ and the Holm-Bonferroni method, to control the familywise type I error at 0.05. Specifically, first, in every permutation, every entry in each measurement was randomly multiplied by −1 or +1, and the t statistic was calculated for the permuted data of each measurement. Next, the maximum t statistic was obtained out of all the measurements. After 5000 permutations, we obtained a surrogate distribution of maximum t statistic, to which we compared the observed t statistic calculated from the actual data in each measurement. The significance level (i.e., p_corrected_) equaled to the proportion of values in the surrogate distribution of maximum t statistic that were greater than the observed t statistic. This permutation procedure was performed iteratively, in that if the measurement with the lowest uncorrected p value survived the test, this measurement was deemed significant after multiple comparisons correction and was excluded from further analysis. Next, the remaining measurements were submitted to the same permutation test again. This procedure was repeated until the measurement with the lowest uncorrected p value did not pass the statistical significance threshold or no measurements were left for testing. Results are depicted in Figure 3a.

For each of the directional one-sample t tests, we calculated the Bayes factor (BF_10_), which indicates the relative likelihood of the alternative hypothesis (i.e., the group mean was greater than 0) over the null hypothesis (i.e., the group mean was not greater than 0) ^68^. The scale r on effect size we adopted was 0.707. BF_10_ greater than 3/10/30 indicates moderate/strong/very-strong evidence for the alternative hypothesis, whereas BF_10_ less than 0.333/0.1/0.03 indicates moderate/strong/very-strong evidence for the null hypothesis ^69^.

To visualize fMRIa, we constructed first-level fMRIa-GLM3, in which different regressors modeled the location occupation periods with different inter-location distances between successively visited locations (i.e., 0m, 4m, 8m, and 12m). We then plotted beta estimates of these regressors (i.e., estimated brain activation levels) as a function of inter-location distance. Results are depicted in Figure 3c.

#### Disentangling objective location and subjective location in fMRIa

We constructed a first-level general linear model (fMRIa-GLM2) to directly compare objective location and subjective location by disentangling their unique contributions to fMRIa. In the model setup, we included two parametric regressors defined by objective location and subjective location in the model, with no orthogonalization. We created two versions of fMRIa-GLM2. The only difference between the two versions was the order in which the parametric regressors were entered into the model. In fMRIa-GLM2a, the objective-location-defined parametric regressor was entered first, followed by the subjective-location-defined parametric regressor. In fMRIa-GLM2b, the order was reversed. We took the beta estimate for the subjective-location-defined parametric regressor in fMRIa-GLM2a as the unique contribution of subjective location, and the beta estimate for the objective-location-defined parametric regressor in fMRIa-GLM2b as the unique contribution of objective location ^70^. Because some participants did not commit any mistakes in some runs, which would result in exactly the same parametric regressors for objective location and subjective location, we concatenated all the scans belonging to the same cue type together across runs and days in SPM12. Run-wise head motions were modeled as nuisance regressors, which resulted in 6 * 16 runs = 96 nuisance regressors in total. Images of the beta estimates for the regressors were left spatially unsmoothed.

At the group-level, in the ROI-based analysis, beta estimates for fMRIa of all the voxels in the ROI were averaged. Then participant-specific beta estimates for the unique contributions of objective location and subjective location were tested using directional one-sample t test, separately.

For each of the directional one-sample t tests conducted here, we calculated the Bayes factor (BF_10_).

Results are depicted in Figure 3b.

#### fMRIa pattern similarity analysis

To investigate the voxel-to-voxel distribution patterns of fMRIa, we developed the fMRIa pattern similarity analysis, which is analogous to the representational similarity analysis (RSA)^71^. RSA is a form of multi-voxel pattern analyses. Conventionally, the multi-voxel pattern analysis is applied to activation levels of voxels in fMRI studies ^72^. Recently, these techniques have been applied to other measurements, e.g., inter-region functional connectivity ^16^. Here, we applied the RSA technique to fMRIa, meaning that the basic elements in the computations were the voxels’ fMRIa magnitudes instead of their activation levels. If the spatially distributed pattern of fMRIa across voxels was distinct between landmarks and self-motion cues, this would indicate that the two cue types recruited dissociable neural representations in terms of fMRIa.

The fMRIa pattern similarity analysis was based on beta estimates of fMRIa as estimated in fMRIa-GLM1. Images of the beta estimates for the regressors were left spatially unsmoothed. The procedure is illustrated in Figure 4a. First, one fMRIa vector was estimated for each run. The fMRIa vector contained the fMRIa estimates of all the voxels in the ROI, with each element of the vector corresponding to the fMRIa magnitude (signed) of a voxel in the ROI. Second, for each cue condition, in each scanning day, we divided the data into two parts based on the chronological order, resulting in four parts in total for each cue type. In this way, for each cue type, each part contained two fMRIa vectors from the two different environments in two consecutive runs. To eliminate any subtle effects of environment, which was not of our primary interest, we computed the mean fMRIa vector by averaging fMRIa for each voxel across the two environments within each part (see a similar treatment to eliminate possible subtle influences of an uninterested factor in fMRI multi-voxel pattern analysis in Shine et al., 2019). This resulted in four mean fMRIa vectors in total for each cue type, with two mean vectors in each scanning day. Third, fMRIa pattern similarity was computed in a cross-validated manner by calculating the Pearson correlation between the mean fMRIa vectors from different parts (Walther et al., 2016). Specifically, within-cue similarity was calculated as the Pearson correlation between the mean fMRI vectors of the same cue type. Within-cue similarity was first calculated for the two cue types separately (i.e., within-landmark similarity and within-motion similarity), and was then averaged across the cue types. Between-cue similarity was calculated in the same manner, but the two mean fMRIa vectors in the correlation calculation were from different cue types. We obtained the final estimates of within-cue similarity and between-cue similarity by averaging all the Pearson correlations (Fisher-transformed) calculated from all possible pairs of the mean fMRIa vectors. Finally, to obtain the fMRIa pattern distinction score, we subtracted between-cue similarity from within-cue similarity. We then tested the fMRIa pattern distinction score against 0, using 1-tailed one sample t tests. A positive fMRIa pattern distinction score would indicate that the voxel-to-voxel spatial distribution of fMRIa was distinct between the two cue types. Importantly, in the analysis, we distinguished between within-day and between-day fMRIa pattern distinction scores, given the possibility that the fMRIa pattern might not necessarily be stable across days within the same cue type.

For each of the t tests conducted in this analysis, we calculated the Bayes factor (BF_10_). Results are depicted in Figure 4b.

#### Analysis of multi-voxel pattern similarity

To analyze multi-voxel pattern similarity of activation vectors (MVPS), we constructed MVPS-GLM1 as the first-level general linear model (GLM), in which separate regressors modeled the location occupation phase for the four test locations. No parametric regressors were included. Other aspects of the model were the same as in the above-mentioned fMRIa-GLM1. The beta images for the regressors were left spatially unsmoothed. Similar to the fMRIa analysis, we created two versions of MVPS-GLM1: in MVPS-GLM1a, the location occupation regressors were defined by objective locations (i.e., where the participant was actually located); in MVPS-GLM1b, the location occupation regressors were defined by subjective locations (i.e., where the participant reported he/she was located).

The MVPS analysis was conducted as follows (Figure 5a). In step 1, for each cue type and scanning day, the dataset was divided into two parts chronologically, resulting in four parts in total for each cue type. For each cue type, within each part, the factor ‘environment’, which was not of our main interest here, was averaged out by computing the mean activation vector of the two consecutive runs belonging to the two environments for each test location. Each element of the mean activation vector denotes the mean activation level averaged across the two runs of each voxel in the ROI. Note that here, the test location was defined by either objective location (MVPS-GLM1a) or subjective location (MVPS-GLM1b), as described above. This resulted in four mean activation vectors for each location and each cue type. In step 2, we calculated cross-validated activation pattern similarities by calculating Pearson correlations between the mean activation vectors of pairwise test locations from different parts. This resulted in the 4×4 activation pattern similarity matrix (Figure 5a.2). In step 3, the activation pattern similarity matrix was averaged element-by-element across all possible part pairs, resulting in the 4×4 mean activation pattern similarity matrix (Figure 5a.3). In step 4, pairwise inter-location distances among the four test locations were calculated, resulting in the 4×4 inter-location distance matrix that contained values of 0m, 4m, 8m, and 12m. In other words, inter-location distance was modeled in a continuous manner. In the final step, the spatial information score was calculated as the Pearson correlation between the mean activation pattern similarity matrix (Fisher-transformed) and the inter-location distance matrix, which was Fisher-transformed and reversed in sign. A positive information score would indicate that spatial distance information among the test locations was encoded in the BOLD signals, meaning that test locations were more similar to each other in neural representations as the distance between them decreased.

We calculated spatial information scores for landmarks, self-motion cues, and between cue types, in which the mean activation vectors in the correlation calculation in step 2 were estimated both from the landmark condition, both from the self-motion condition, and from different cue conditions, respectively. Importantly, the between-cue spatial information score would be informative of whether the neural coding of spatial distance information was generalizable between different cue types. We calculated spatial information scores based on objective location using MVPS-GLM1a or subjective location using MVPS-GLM1b.

At the group-level, the six measurements of spatial information scores (i.e., location type (objective-location-based vs. subjective-location-based) X measurement type (landmark vs. self-motion vs. between-cue)) were tested using directional one-sample t tests separately, to obtain the uncorrected significance levels (i.e., p_uncorrected_). Then, to control the familywise type I error at 0.05, the six measurements were submitted to a multiple comparisons correction approach that combines the nonparametric permutation-based maximum-t-statistic method ^67^ and the Holm-Bonferroni method, as described in the previous section on fMRIa.

For each of the directional one-sample t tests conducted here, we calculated the Bayes factor (BF_10_).

Results are depicted in Figure 5b and Figure 5d.

#### Disentangling objective location and subjective location in MVPS

To directly compare objective location and subjective location in MVPS, we attempted to estimate the unique contributions of objective and subjective location to the overall MVPS by conducting the following analysis. In the first-level general linear model MVPS-GLM2, we modeled individual trials with separate regressors. Each trial was associated with two location labels, one defined by objective location and the other defined by subjective location. Whether the two labels matched or mismatched depended on the behavioral accuracy in that trial. We computed cross-validated Pearson r correlation between single-trial-based activation patterns from two different runs, resulting in a 20×20 activation pattern similarity matrix for a run pair. Two 20×20 inter-location distance matrices were constructed, one based on objective location and the other on subjective location. We then used these two inter-location distance matrices (standardized) to predict the 20×20 activation pattern similarity matrix (fisher-transformed and standardized) using the multiple linear regression analysis for each run pair. The two regression coefficients (i.e., beta-unique; reversed in sign) denoted the respective unique contributions of the two predictors, with the contributions of the other predictor excluded. The multiple linear regression was performed for each run pair, and the estimated regression coefficients were then averaged across all run pairs to obtain the final estimates of unique contributions of objective location and subjective location, which were then tested against 0 using directional one-sample t tests. Bayes factors (BF_10_) were also computed.

This analysis was conducted for the landmark condition, self-motion condition, and between cue types, separately. For the landmark condition and self-motion condition, the two runs in each run pair were from the same cue type, whereas for between cue types, they were from different cue types. Run pairs were assigned with a value of ‘NaN’ for the beta-unique estimates and excluded from further analysis, when the behavioral performance was perfect in both runs (i.e., the objective-location-based and the subjective-location-based inter-location distance matrices were identical and perfectly correlated with each other). When all the participants were considered, for the landmark condition, 64 out of 28 * 20 subjects = 560 run pairs (= 11.43%) had 100% accuracy rate and ‘NaN’ as the beta-unique estimates. For the self-motion condition, 18 out of 560 run pairs (= 3.21%) had 100% accuracy rate. For between cue types, 31 out of 1280 run pairs (= 2.42%) had 100% accuracy rate).

Results are depicted in Figure 5c.

#### Neural space reconstruction analysis

As an overview, in the neural space reconstruction analysis (Figure 6a), first, a certain form of neural distance matrix was constructed for objective test locations, depending on the type of fMRI effect being investigated. Elements in the neural distance matrix denote pairwise neural distances between the test locations. Next, multi-dimensional scaling was performed on the neural distance matrix to recover the spatial coordinates of the locations in the neural space, following the basic principle that locations with greater representational similarities are positioned closer to each other in the neural space ^74^. Finally, the Procrustes analysis was performed to map the estimated coordinates of the locations to the original physical space through rotations and reflections ^75^. The neural space reconstruction analysis is commonly applied to multi-voxel activation patterns in fMRI studies ^76^. In the current study, we applied this analysis to fMRIa, in addition to MVPS (see the rationale below). The neural space was reconstructed based on the neural distances between objective test locations (instead of participants’ subjective locations).

The procedure was the same for both MVPS and fMRIa, except for how the neural distance matrix was constructed. For MVPS, in step 1, the neural distance between two test locations (defined by objective location) was quantified by the degree of correlational dissimilarity between their voxel-to-voxel activation patterns (e.g., 1-Pearson correlation), based on MVPS-GLM1a. The neural distance between two test locations indicated how ‘dissimilar’ they were in neural representations. To obtain the neural distance matrix, we constructed the 4×4 representational dissimilarity matrix, with each element equal to 1 minus the Pearson correlation between the activation patterns of two test locations. In step 2, we averaged symmetrical off-diagonal elements in the matrix. The four diagonal entries were manually set to 0, because multidimensional scaling only exploits relative distances between different items (also see ^77^). Elements in the matrix were normalized to be within the range [0, 1] as follows: normalized value= (original value-matrix minimum)/(matrix maximum – matrix minimum). In step 3, the normalized neural distance matrix was subjected to multidimensional scaling and the Procrustes analysis.

For fMRIa, the neural distance between two test locations (defined by objective location) could be quantified as the brain region’s activation level for one location when preceded by the other location - the lower the region’s activation to the current location when preceded by the other location, the larger the repetition suppression effect, the closer the two test locations would be positioned to each other in the neural space. In step 1, we constructed the adaptation matrix, which is parallel to the representational dissimilarity matrix in MVPS. We relied on MVPS-GLM2, which modeled the location occupation phase in individual single trials with separate regressors. These trials were classified into 4**×**4 = 16 groups based on the combination of two locations visited in succession; the beta estimates for trials from the same group were averaged, resulting in the 4×4 adaptation matrix. In the adaptation matrix, rows represent the previous location, columns represent the current location, and each element represents the activation level at the current location when preceded by the previous location. To keep it consistent with the main fMRIa analysis, the fMRIa-based neural space reconstruction analysis was restricted to trials that could be modeled for fMRIa (i.e., locations not preceded by the null event and not the first event in the sequence).

Nevertheless, to confirm that the baseline activation level was comparable for the four test locations in RSC and the hippocampus, we estimated activation levels of RSC and hippocampus for the four test locations using the trials that were not included in the parametric regressors modeling fMRIa (i.e., test locations preceded by the null event and the first event in the sequence in fMRIa-GLM1). We observed no significant differences among the four locations in either RSC (F(3,57) = 0.742, p = 0.531, η_p_^2^ = 0.038) or hippocampus (F(3,57) = 0.875, p = 0.459, η_p_^2^ = 0.044). In addition, there were no significant differences among the four locations in baseline activation in any other ROIs in the medial temporal lobe (Fs <2.1, ps > 0.1, η_p_^2^ < 0.1). This verifies our choice of using the estimated brain activation level for the current location as an indicator of the neural distance between the current location and the preceding location in fMRIa.

The following two steps were the same as in the MVPS-based neural space reconstruction analysis. In step 2, we normalized the adaptation matrix as to render all the 16 elements within the range [0, 1]. Elements that were diagonally symmetrical to each other in the matrix were averaged, and the diagonal elements were manually set to 0. In step 3, the normalized neural distance matrix was then subjected to multidimensional scaling and the Procrustes analysis.

To address the question of whether the neural space resembled the behavioral performance pattern), we performed the neural space reconstruction analysis for each participant. The reconstructed distances were submitted to a repeated-measures ANOVA test, and with cue type (landmark vs. self-motion) and adjacent location pair (Loc1-2, Loc2-3, Loc3-4) as independent variables. We were particularly interested in the interaction effect between cue type and the linear trend of adjacent location pair, motivated by the observation of differential representational precision patterns between the two cue types (Figure 2). Results are depicted in Figure 6b.1 & 6c.1.

We also addressed the question of whether the neural space resembled the original physical space. To increase statistical power, the normalized neural distance matrix was averaged across participants to obtain the grand group-level neural distance matrix for the four test locations (defined by objective location) ^77–79^, which was then subjected to multidimensional scaling and the Procrustes analysis. We performed a nonparametric permutation test as follows. First, we obtained the actual Procrustes distance calculated from the group-level neural distance matrix. Procrustes distance indicates the deviation of the reconstructed neural space from the original physical space. Second, we applied the permutation procedure to obtain the surrogate distribution of Procrustes distance, to which the actual Procrustes distance would be compared. Specifically, in each permutation, we randomly shuffled the 12 off-diagonal entries in the grand group-level neural distance matrix. Note that to allow for more permutations, this shuffling was done prior to the averaging of symmetrical off-diagonal elements in the neural distance matrix. We obtained the Procrustes distance by applying multidimensional scaling and the Procrustes analysis to the shuffled neural distance matrix. This process was repeated 5000 times, resulting in a surrogate distribution of Procrustes distance. Third, the actual Procrustes distance was compared to the surrogate distribution. The significance level (i.e., p value) was calculated as the proportion of values in the surrogate distribution being smaller than the actual Procrustes distance, analogous to directional one-sample t test. Results are depicted in Figure 6b.2 & 6c.2.

#### Assessing spatial overlap between MVPS and fMRIa at the voxel level

To assess the spatial overlap between fMRIa and MVPS, we performed a fMRIa-based artificial lesion analysis, in which we selectively excluded a certain proportion of voxels based on their fMRIa magnitudes (spatially unsmoothed) prior to calculating the spatial information score in MVPS ^80, 81^. Since our previous results showed that in RSC, fMRIa was mainly objective-location-driven and MVPS effect was mainly subjective-location-driven, we conducted this analysis using objective-location-based fMRIa as estimated from fMRIa-GLM1a and subjective-location-based MVPS as calculated from MVPS-GLM1b. We ranked voxels in the ROI by the landmark or self-motion fMRIa magnitude (signed) from low to high, using the unsmoothed beta images estimated from fMRIa-GLM1a. We conducted the MVPS analysis (Figure 5a) with one quarter of voxels excluded at one time. To address the question of whether voxels’ fMRIa levels affected MVPS, we conducted a repeated-measure ANOVA test, with the excluded quarter as the independent variable and the resulted spatial information score as the dependent variable.

As a critical comparison, we calculated the empirical chance level of the resulted spatial information score, by conducting the same artificial lesion analysis, but with the voxels randomized in order instead of being ordered by the fMRIa magnitude. In each randomization, we calculated the spatial information score after deleting one quarter of the voxels. Voxel randomization was performed for 1000 times, and the mean resulted spatial information score averaged across all the randomizations was taken as the empirical chance level. Hence, the relative contribution of a certain voxel group can also be assessed by comparing the resulted spatial information score to the empirical chance level.

Results are depicted in Figure S7.

#### Analysis of empirical relative detection power for fMRI adaptation

In fMRI data analysis, blood-oxygen-level-dependent (BOLD) signals of certain frequencies are attenuated or even eliminated: first, the convolution with the hemodynamic response function (HRF) dampens high-frequency signals; second, the high-pass filter eliminates low-frequency signals. This means that only a proportion of the original BOLD signals will be retained in further analysis, which is termed as ‘relative detection power (*DP_rel_*,). Specifically, *DP_rel_*, is calculated as follows ^19^,

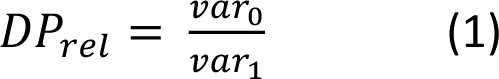

in which *var*_0_ represents the hypothesized neural modulation after HRF convolution and high-pass filtering (e.g., f > 1/128), and *var*_1_ represents the original hypothesized neural modulation prior to HRF convolution and high-pass filtering. *DP_rel_* ranges from 0 to 1. To interpret, *DP_rel_* of one means no loss of detection power, and *DP_rel_* of zero means a complete loss. Therefore, *DP_rel_* reflects the probability for us to detect effects in fMRI BOLD signals.

In the current study, we used the eight de Bruijn sequences from our previous study ^3^. These sequences were generated based on objective locations, and hence, were theoretically optimized in terms of *DP_rel_* with respect to objective locations. However, in the current study, participants’ responses could not be known in advance, leading to the possibility that *DP_rel_* was reduced for the subjective-location sequences compared to the objective-location sequences. Therefore, our observation that fMRI adaptation was predominantly driven by objective location rather than subjective location could have been confounded by potentially higher *DP_rel_* for the objective-location sequences than the subjective-location sequences.

To address this issue, we calculated the empirical *DP_rel_* for objective-location and subjective-location sequences separately, based on the first-level general linear models (GLMs) using events and inter-location distances that actually occurred in the experiment for each participant. These GLMs included regular regressors modeling the location occupation events as a measure of the direct stimulus effect, and parametric regressors modeling the inter-location distance as a measure of the adaptation effect, same as in the construction of fMRIa-GLM1a and fMRIa-GLM1b in the main analysis.

Specifically, to calculate *var*_1_ in equation (1), we constructed these first-level GLMs with no HRF convolution and no high-pass filtering applied. We then converted the simulated BOLD signal of the parametric regressor from the time domain to the frequency domain, using the fast Fourier transform (FFT). The variance of the hypothesized neural modulation for the parametric regressor (i.e., *var*_1_) was calculated as the area under curve (AUC) using the frequency-domain data.

To calculate *var*_0_ in equation (1), we convolved the predicted fMRI time-series for the parametric regressor with the canonical hemodynamic response function (HRF), and adopted a high-pass filter with a cut-off at 1/128s = 0.0078 Hz. The variance of the convolved and filtered signal for the parametric regressor (i.e., *var*_0_) was calculated in the same way as *var*_1_. Finally, to obtain *DP_rel_* we divided *var*_0_ over *var*_1_.

## Videos

**Title: Demo of the experimental environments and tasks, related to** **Figure 1** **and the section ‘Stimuli and navigation task’ in STAR Methods.** Demo of the learning trials starts at 0’0” and ends at 1’24”. Demo of the location identification task (i.e., test) starts at 1‘25“and ends at 2’48“.

Video file: Learning_and_location_identification_task_demo.mp4

---

**Figure S1.**
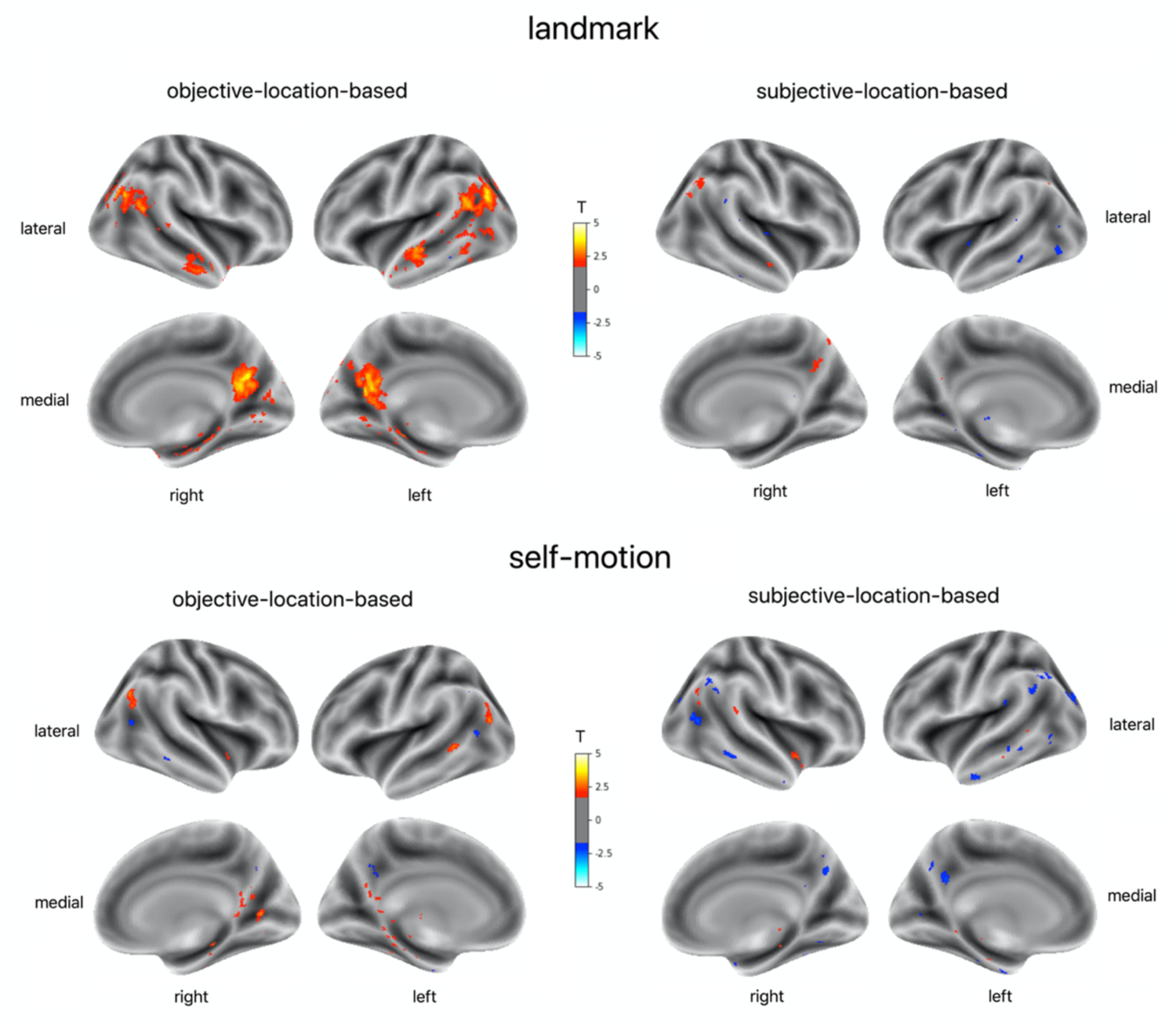
Voxel-wise analysis of fMRIa in the entire volume, related to Figure 3. Results are displayed for the landmark condition (upper) and the self-motion condition (lower), objective-location-based fMRIa (left) and subjective-location-based fMRIa (right). The parametric regressors modeled same vs. different locations in the landmark condition, and continuous inter-location distance in the self-motion condition, as in the main analyses (Figure 3). The participant-specific maps of fMRIa were normalized to the MNI template and spatially smoothed with 3mm isotropic FWHM. For the 2^nd^ level analysis, we conducted directional one-sample t test against 0. The parametric t maps were overlaid on the MNI template and projected to the brain surface. Here, results are thresholded at p_uncorrected_ = 0.05. When corrected for multiple comparisons across the entire volume using the nonparametric permutation test (Nichols & Holmes, 2002), there were no significant voxels with the voxel-inference approach. When the cluster-inference approach (voxel-wise t > 3) was adopted, in the landmark objective location condition, there were three significant clusters (p_FWE-corr_ < 0.05, 1-tailed), encompassing the angular gyrus (MNI coordinates of local maxima: [50, −71, 30], [−45, −71, 28]), middle occipital gyrus (MNI coordinates of local maxima: [40, −79, 35], [45, −77, 27], [−34, −84, 33]), calcarine (MNI coordinates of local maxima: [3, −56, 12], [−12, −46, 7]), and precuneus (MNI coordinates of local maxima: [1, −63, 24]). In the other conditions, no significant clusters were detected.

**Figure S2.**
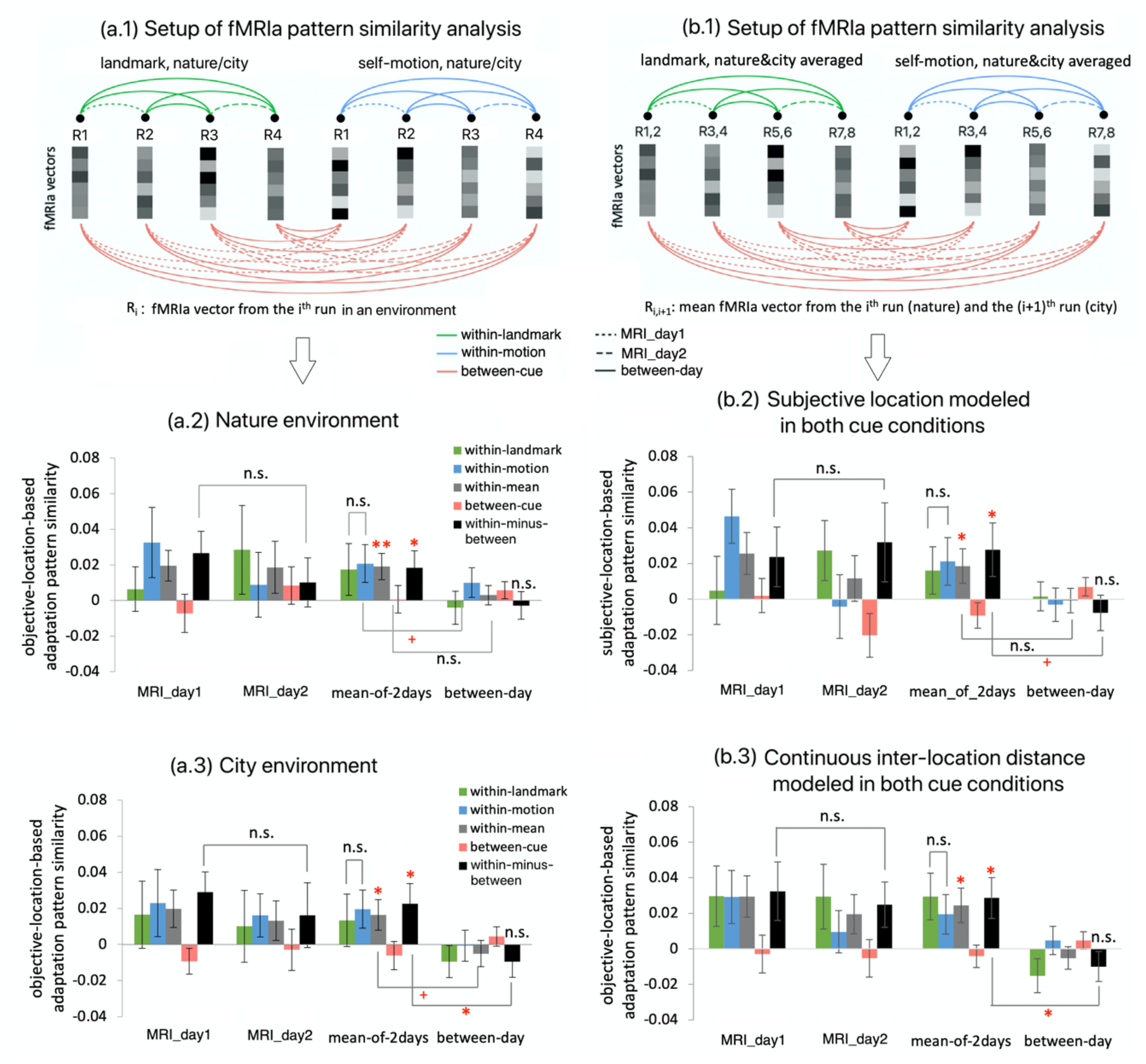
Controlled analyses of fMRIa pattern similarity in retrosplenial cortex, related to Figure 4 and the main text section ‘fMRIa-based distance coding was spatially distinct between cue types in RSC’. (a) Separate analyses of objective-location-based fMRIa pattern similarity in the nature environment (a.2) and the city environment (a.3), with inter-location distance modeled continuously in the self-motion condition and ‘same vs. different locations’ modeled in the landmark condition, as in the main analysis (Figure 4). (a.1) Setup of the analyses is the same as in the main analysis (Figure 4a), except that fMRIa vectors were not averaged across different environments. (b) More controlled analyses. (b.2) Controlled analysis on subjective-location-based pattern similarity, with inter-location distance modeled continuously in the self-motion condition and ‘same vs. different locations’ modeled in the landmark condition, as in the main analysis (Figure 4). (b.3) Controlled analysis on objective-location-based fMRIa pattern similarity with inter-distance modeled continuously in both cue conditions. (b.1) Setup of the analyses is the same as in the main analysis (Figure 4a). All the controlled analyses revealed a pattern of results similar to the main analysis (Figure 4b). n.s. denotes p_1-tailed/2-tailed_ > 0.1; + denotes p_1-tailed/2-tailed_ < 0.1; * denotes p_1-tailed/2-tailed_ < 0.05; ** denotes p_1-tailed/2-tailed_ < 0.01. Error bars represent ± S.E..

**Figure S3.**
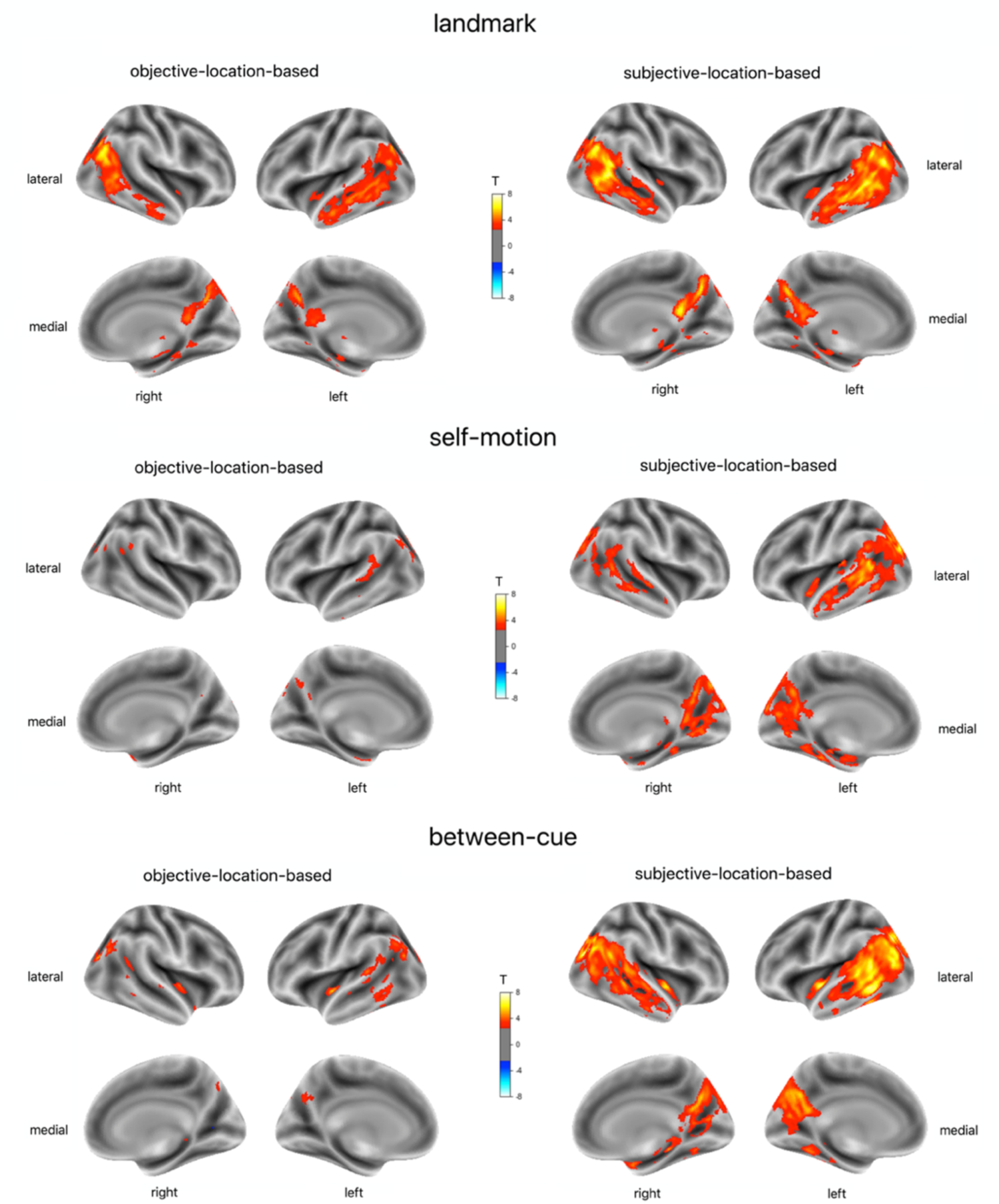
Searchlight analysis of MVPS in the entire volume, related to Figure 5. Results are displayed for the landmark condition (upper), self-motion condition (middle), and between cue types (lower), and for objective-location-based MVPS (left), and subjective-location-based MVPS (right). In all situations, the inter-location distance was modeled continuously, with distances of 0m, 4m, 8m, and 12m, as in the main analysis (Figure 5). The searchlight analysis was conducted in each participant’s native brain, using codes adapted from the TDT toolbox (Hebart et al., 2015) and a searchlight radius of 6mm. At each step, for voxels within the searchlight, the spatial information score was calculated, using the same procedure shown in Figure 5a; the score was then assigned to the voxel at the center of the searchlight. The participant-specific brain maps of spatial information score were normalized to the MNI template and spatially smoothed with 3mm isotropic FWHM. For the 2^nd^ level analysis, we conducted directional one-sample t test against 0. Here, the parametric t maps were overlaid on the MNI template and projected to the brain surface. Results are thresholded at p_uncorrected_ < 0.01. Detailed results are listed in Table S2.

**Figure S4.**
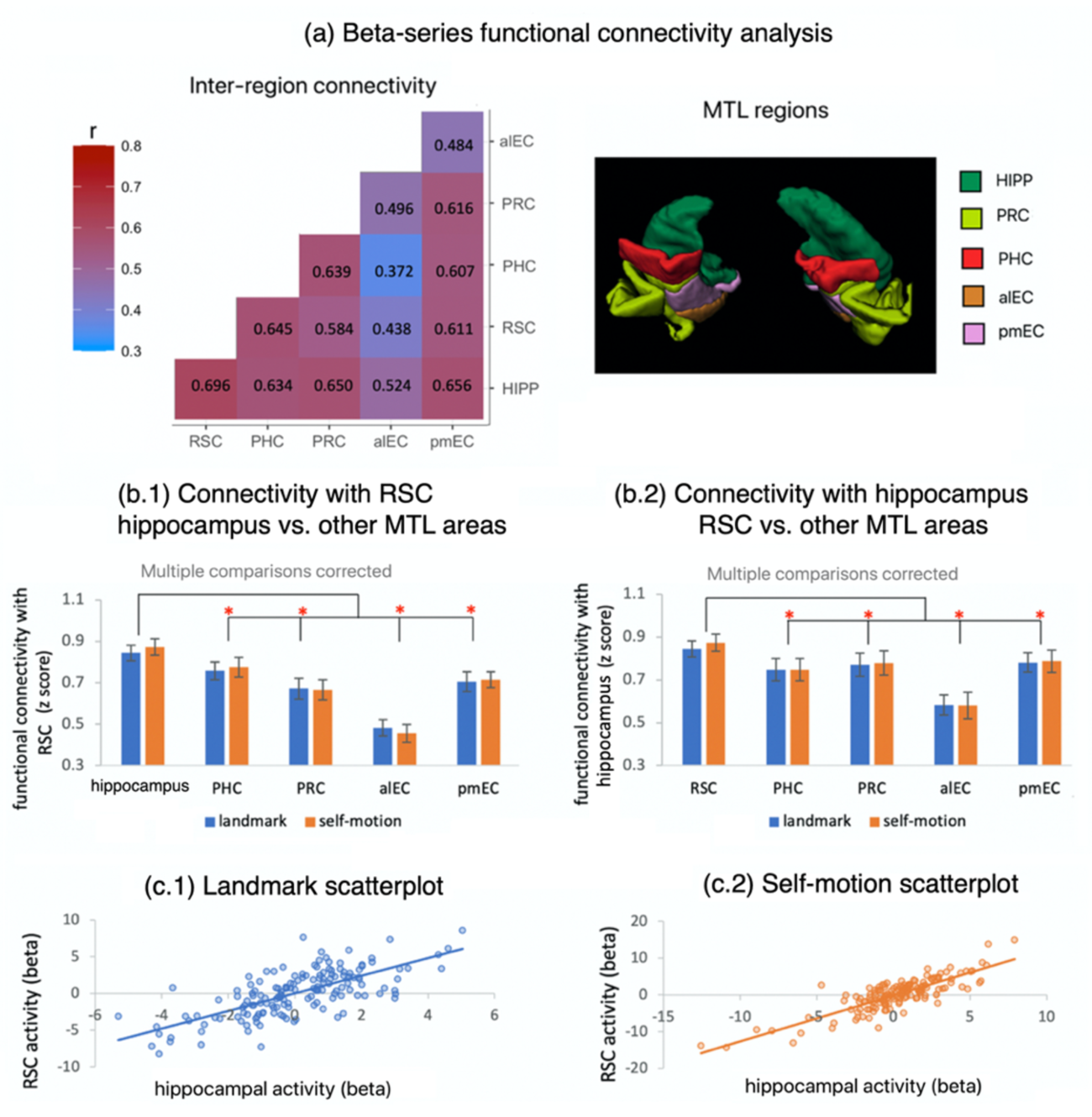
Functional connectivity between retrosplenial cortex and hippocampus, related to the main text section ‘Hippocampus contained a spatial coding scheme similar to RSC’. (a) Results of the beta-series functional connectivity analysis. Displayed on the left is the mean pairwise simple correlations among RSC and the medial temporal lobe (MTL) regions. Displayed on the right are anatomical masks of MTL regions for an exemplary participant. We assessed the functional connectivity between these regions using the beta-series connectivity analysis (Cisler et al., 2014), using MVPS-GLM2 that modeled individual trials with separate regressors (STAR Methods). For each brain region, we obtained a temporal sequence of activation estimates concatenated across individual trials, which were mean-centered within each run prior to the trial concatenation. We then calculated pairwise Pearson r correlations (fisher-transformed) between the temporal sequences of these regions for each participant. There existed strong functional coupling between RSC and hippocampus in both the landmark condition (p_2-tailed_ < 0.001, BF_10_ > 1000) and the self-motion condition (p_2-tailed_ < 0.001, BF_10_ > 1000). (b.1) The five pairs differed significantly in connectivity (F(4,76) = 36.079, p < 0.001, η_p_^2^= 0.655). Planned comparisons showed that RSC-hippocampus connectivity was significantly stronger than RSC’s connectivity with other MTL regions. (b.2) The five pairs differed significantly in connectivity (F(4,76) = 15.549, p < 0.001, η_p_^2^ = 0.450). Planned comparisons showed that RSC-hippocampus connectivity was significantly stronger than the hippocampus’s connectivity with other MTL regions. Effects involving cue type were not significant. * denotes p_holm,2-tailed_ < 0.05. (c) Scatterplot of trial-by-trial activation of the hippocampus and RSC in the landmark condition (c.1) and the self-motion condition (c.2) in an exemplary participant. RSC: retrosplenial cortex; HIPP: hippocampus; PHC: parahippocampal cortex; PRC: perirhinal cortex; alEC: anterior-lateral entorhinal cortex; pmEC: posterior-medial entorhinal cortex.

**Figure S5.**
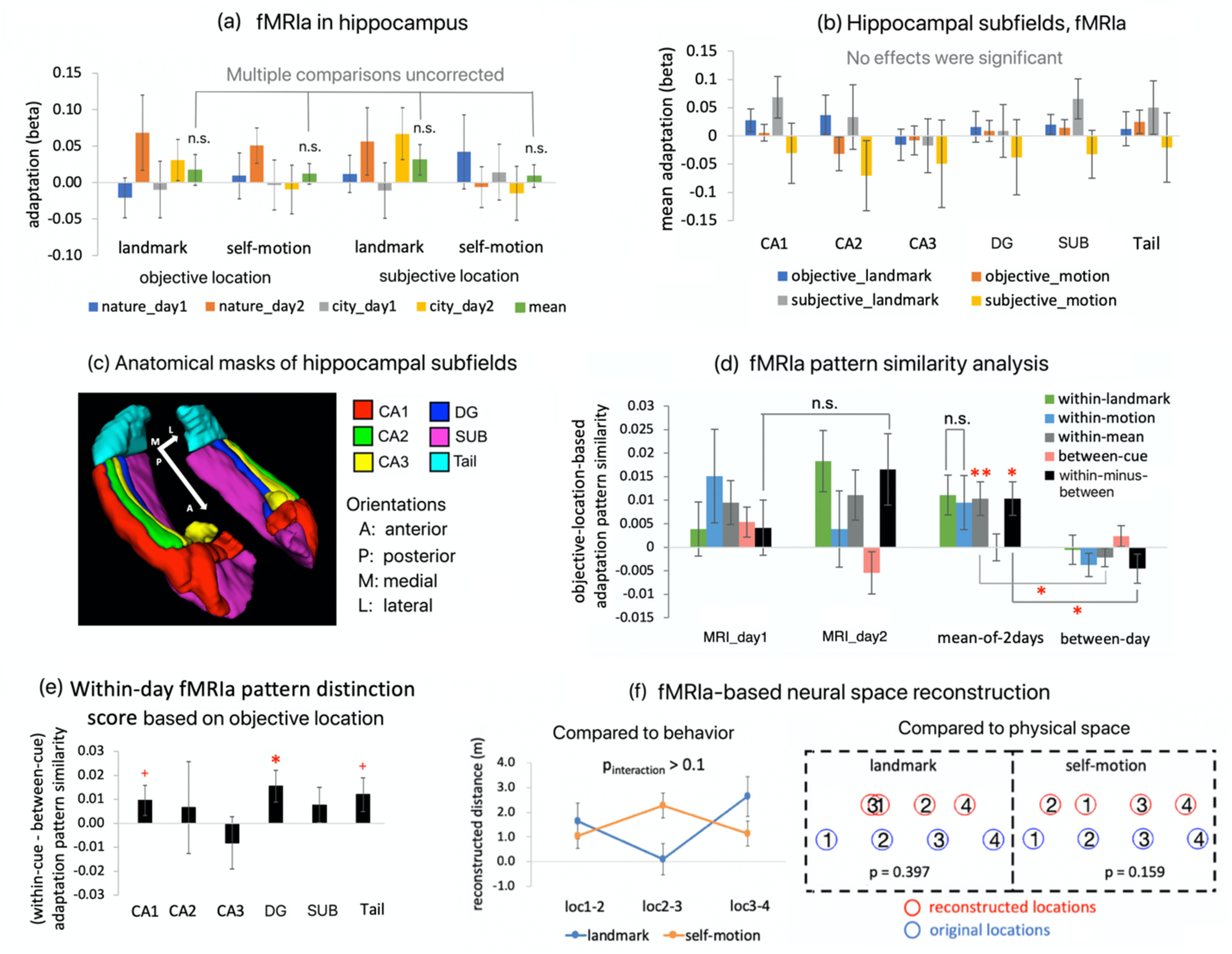
fMRIa results in hippocampus, related to the section ‘Hippocampus contained a spatial coding scheme similar to RSC’ in the main text and Figure 3&4&6. (a) The univariate fMRIa analysis that assessed objective-location-based and subjective-location-based fMRIa for landmarks and self-motion cues, displayed separately for different environments and scanning days. The continuous inter-location was modeled for both cue types by default, because a previous study reporting fMRIa-based neural coding of continuous distance between locations defined by landmarks in the hippocampus (Morgan et al., 2011). The mean fMRIa (green bars) was not significant for either cue type, even at the uncorrected significance level. (b) Mean fMRIa averaged across environments and days is displayed for each hippocampal subfield. No significant fMRIa was observed in any subfields. (c) Anatomical masks of hippocampal subfields for an exemplary participant (DG – dentate gyrus; SUB - subiculum). (d) fMRIa pattern similarity analysis based on objective location. Setup of this analysis is identical to Figure 4a. The within-day fMRIa pattern distinction score was significantly positive (t(19) = 2.090, p_1-tailed_ = 0.018, BF_10_ = 2.682). This implies potential spatial coding in the hippocampus, though fMRIa averaged across voxels was not significant in the hippocampus (a). (e) Within-day fMRIa pattern distinction score for each hippocampal subfield. The score reached statistical significance in the dentate gyrus (DG, t(19) = 1.930, p_1-tailed_ = 0.034, BF_10_ = 2.082), but not in other subfields (ps > 0.05). (f) Results of the neural space reconstruction analysis based on fMRIa in the hippocampus. The reconstructed neural spaces did not resemble participants’ behavior (left) or the original physical space in any cue conditions (right). n.s. denotes p_1-tailed/2-tailed_ > 0.1, * denotes p_1-tailed/2-tailed_ < 0.05, and ** denotes p_1-tailed/2-tailed_ < 0.01; + denotes p_1-tailed/2-tailed_ < 0.1.

**Figure S6.**
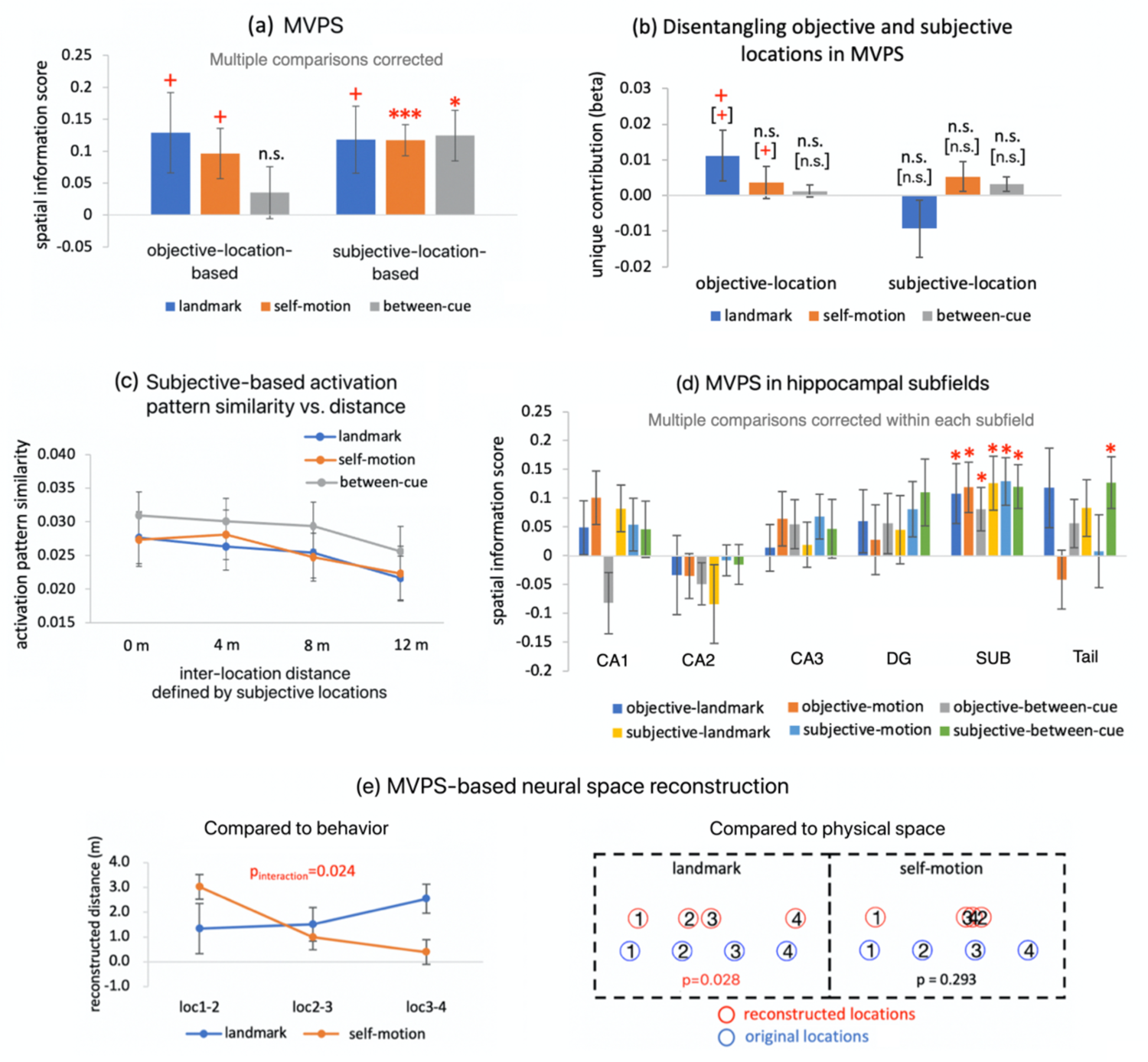
MVPS results in the hippocampus, related to section ‘Hippocampus contained a spatial coding scheme similar to retrosplenial cortex’ in the main text and to Figure 5&6. (a) Spatial information score based on objective location and subjective location for landmarks, self-motion cues, and between cue types. The spatial information score was significant or marginally significant for all the three measurements (landmark, self-motion, between-cue) when based on subjective location. (b) Unique contributions of objective location or subjective location were not significant for any cue type or location type (objective vs. subjective). (c) To visualize the MVPS effects, activation pattern similarity is plotted as a function of inter-location distance defined by subjective location for landmarks, self-motion cues, and between cue types. (d) MVPS results in each hippocampal subfield. Statistical results were corrected for multiple comparisons within each subfield using the nonparametric permutation test and the Holm-Bonferroni procedure (STAR Methods). In the subiculum (SUB), spatial information score was significant for all three measurements (landmark, self-motion, and between-cue) and for both location types (objective and subjective location). DG - dentate gyrus. (e) In the hippocampus, the MVPS-based neural spaces significantly resembled the participants’ behavior (i.e., the interaction between cue type and the linear trend of location pair was significant, p_interaction_ = 0.024), resembled the physical space in the landmark condition (p = 0.028), but did not resemble the physical space in the self-motion condition (p = 0.293). * denotes p_1-tailed_ < 0.05, and + denotes p_1-tailed_ < 0.1, n.s. denotes p_1-tailed_ > 0.1, * denotes p_1-tailed_ < 0.05, and ** denotes p_1-tailed_ < 0.01; + denotes p_1-tailed_ < 0.1.

**Figure S7.**
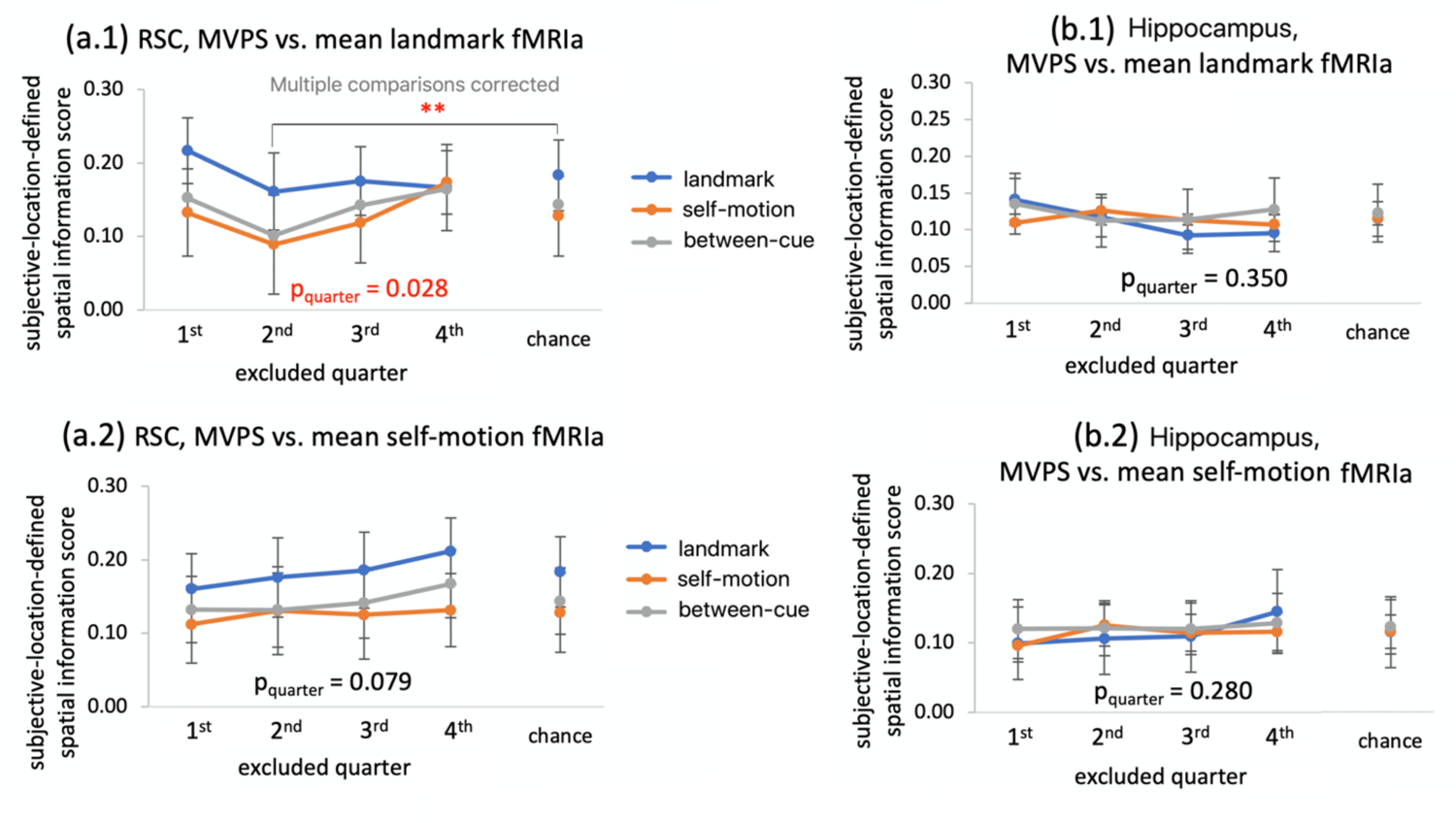
Results of the fMRIa-based artificial lesion analysis, related to the discussion section in the main text. We conducted these analyses with objective-location-based fMRIa and subjective-location-based MVPS. (a.1-a.2) Results of the fMRIa-based artificial lesion analysis for the retrosplenial cortex (RSC) when voxels were ranked from low to high by mean landmark fMRIa (a.1) and mean self-motion fMRIa (a.2). (b.1-b.2) Results of the fMRIa-based artificial lesion analysis for the hippocampus when voxels were ranked by mean landmark fMRIa (b.1) and mean self-motion fMRIa (b.2). In (a.2) (b.1) and (b.2), the main effect of excluded quarter was not significant (ps < 0.05). The main effect of excluded quarter was significant in (a.1) when the voxels were ranked by mean fMRIa for landmarks (F(3,57) = 4.119, p_quarter_ = 0.028, η_p_^2^ = 0.178): excluding voxels relatively lower in mean landmark fMRIa (i.e., the 2^nd^ quarter) tended to result in lower spatial information scores than excluding other quarters of voxels. Excluding the 2^nd^ quarter of voxels also resulted in significant lower spatial information scores than the empirical chance levels. Details of these analyses can be found in STAR Methods. * denotes p_2-tailed_ < 0.05.

**Table S1.** Influences of environment on univariate fMRIa and MVPS in retrosplenial cortex, related to Figure 3a and Figure 5b. (a) fMRIa is summarized as a function of location type (objective vs. subjective), cue type (landmark vs. self-motion), and environment (nature vs. city). fMRIa did not differ between the two environments (ps > 0.2). (b) Spatial information score in MVPS is summarized as a function of location type (objective vs. subjective), cue type (landmark vs. self-motion vs. between-cue), and environment type (nature vs. city vs. between-city). Similar to the factor day (Figure 5b), environment did not significantly modulate spatial information score (ps > 0.05): the within-nature scores did not differ from the within-city scores, and the within-environment scores (mean of the within-nature and within-motion scores) did not differ from the between-environment scores. Furthermore, also similar to the factor day (Figure 5b), the between-environment spatial information score was significant for all three measurements (landmark, self-motion, and between-cue) based on subjective location, indicating that the spatial coding was generalized between different environments. n.s., p > 0.1; * p < 0.05; ** p < 0.01; *** p< 0.001.

| fMRIa |  |  |  |  |  |
| --- | --- | --- | --- | --- | --- |
| Cue | Environment | Objective-location-based |  | Subjective-location-based |  |
|  |  | Mean | Std. Deviation | Mean | Std. Deviation |
| landmark | nature | 0.108 | 0.257 | 0.086 | 0.232 |
|  | city | 0.089 | 0.160 | 0.061 | 0.175 |
| self-motion | nature | 0.166 | 0.205 | 0.092 | 0.235 |
|  | city | 0.006 | 0.267 | 0.021 | 0.261 |
| MVPS (spatial information score) |  |  |  |  |  |
| Cue | Environment | Objective-location-based |  | Subjective-location-based |  |
|  |  | Mean | Std. Deviation | Mean | Std. Deviation |
| landmark | within-nature | 0.118 | 0.278 | 0.016 | 0.023 |
|  | within-city | 0.128 | 0.255 | 0.018 | 0.032 |
|  | between environment | 0.146 ** | 0.226 | 0.012 ** | 0.018 |
| self-motion | within-nature | 0.049 | 0.236 | 0.005 | 0.031 |
|  | within-city | 0.010 | 0.227 | 0.004 | 0.026 |
|  | between environment | 0.095 * | 0.192 | 0.011 *** | 0.012 |
| between cue | within-nature | 0.125 | 0.220 | 0.007 | 0.019 |
|  | within-city | -0.006 | 0.283 | 0.004 | 0.020 |
|  | between environment | 0.033 n.s. | 0.152 | 0.006 ** | 0.008 |

**Table S2.** The searchlight analysis of MVPS, corresponding to Figure S2 and related to Figure 5. Listed are region (AAL atlas), MNI coordinates (x, y, z), t-value, corrected p-value of the peak voxel (p_FWE-corrected_, 1-tailed, voxel-inference, nonparametric permutation test, Nichols & Holmes, 2002), and cluster size at p_FWE-corrected_ = 0.05. Mid: middle; Inf: inferior; Sup: superior.

| AAL region | Left hemisphere |  |  | Right hemisphere |  |  | T value | pFWE-corr | Cluster size |
| --- | --- | --- | --- | --- | --- | --- | --- | --- | --- |
|  | x | y | z | x | y | z |  |  |  |
| landmark, objective-location-based |  |  |  |  |  |  |  |  |  |
| Angular |  |  |  | 43 | -68 | 31 | 7.76 | 0.004 | 810 |
| Temporal_Mid |  |  |  | 42 | -61 | 22 | 7.36 | 0.007 |  |
| Temporal_Mid | -60 | -46 | -3 |  |  |  | 6.88 | 0.013 | 48 |
| Precuneus |  |  |  | 3 | -65 | 31 | 6.54 | 0.024 | 21 |
| Occipital_Mid | -34 | -75 | 35 |  |  |  | 7.14 | 0.009 | 179 |
|  |  |  |  | 34 | -74 | 38 | 6.42 | 0.028 | 21 |
| landmark, subjective-location-based |  |  |  |  |  |  |  |  |  |
| Temporal_Mid |  |  |  | 58 | -63 | 7 | 9.34 | <0.001 | 718 |
| Temporal_Mid | -62 | -47 | -2 |  |  |  | 8.94 | 0.001 | 1238 |
|  | -57 | -53 | 3 |  |  |  | 7.56 | 0.003 |  |
| Temporal_Mid |  |  |  | 45 | -52 | 16 | 8.51 | 0.001 | 370 |
|  |  |  |  | 50 | -57 | 20 | 6.36 | 0.027 |  |
| Temporal_Mid | -50 | -64 | 21 |  |  |  | 8.47 | 0.001 | 1236 |
|  | -54 | -34 | -8 |  |  |  | 7.10 | 0.007 | 68 |
|  |  |  |  | 47 | -63 | 1 | 6.67 | 0.015 | 79 |
|  | -60 | -28 | -7 |  |  |  | 6.47 | 0.021 | 25 |
|  | -47 | -63 | 0 |  |  |  | 6.26 | 0.037 | 11 |
|  |  |  |  | 57 | -51 | 15 | 6.09 | 0.043 | 6 |
| Occipital_Mid |  |  |  | 45 | -67 | 29 | 8.01 | 0.001 | 571 |
|  |  |  |  | 35 | -75 | 40 | 6.45 | 0.022 | 22 |
|  | -34 | -76 | 40 |  |  |  | 6.27 | 0.031 | 36 |
|  |  |  |  | 39 | -67 | 7 | 6.09 | 0.044 | 2 |
| Precuneus |  |  |  | 3 | -67 | 34 | 7.63 | 0.003 | 254 |
|  |  |  |  | 7 | -46 | 17 | 7.63 | 0.003 | 179 |
| Temporal_Inf | -54 | -57 | -8 |  |  |  | 6.17 | 0.038 | 11 |
| self-motion, objective-location-based |  |  |  |  |  |  |  |  |  |
| Hippocampus | -21 | -11 | -22 |  |  |  | 8.26 | 0.001 | 19 |
| Occipital_Sup | -27 | -84 | 39 |  |  |  | 6.31 | 0.039 | 3 |
| self-motion, subjective-location-based |  |  |  |  |  |  |  |  |  |
| Occipital_Mid | -28 | -79 | 17 |  |  |  | 8.31 | 0.003 | 395 |
| undefined | -18 | -79 | 16 |  |  |  | 6.51 | 0.029 |  |
| Occipital_Sup | -25 | -85 | 39 |  |  |  | 8.05 | 0.005 | 186 |
| undefined |  |  |  | 33 | -62 | 7 | 6.8 | 0.020 | 12 |
| Temporal_Mid | -49 | -50 | 10 |  |  |  | 6.68 | 0.023 | 30 |
|  | -57 | -44 | 0 |  |  |  | 6.29 | 0.043 | 5 |
| Occipital_Mid | -34 | -86 | 24 |  |  |  | 6.39 | 0.036 | 12 |
|  | -32 | -69 | 34 |  |  |  | 6.22 | 0.047 | 2 |
| Cuneus |  |  |  | 12 | -78 | 19 | 6.24 | 0.045 | 1 |
| between-cue, objective-location-based |  |  |  |  |  |  |  |  |  |
| undefined | -45 | -15 | -11 |  |  |  | 8.93 | 0.001 | 78 |
| Occipital_Mid |  |  |  | 33 | -83 | 20 | 7.34 | 0.006 | 49 |
| between-cue, subjective-location-based |  |  |  |  |  |  |  |  |  |
| Occipital_Mid |  |  |  | 32 | -83 | 20 | 10.55 | <0.001 | 509 |
|  |  |  |  | 36 | -76 | 25 | 6.02 | 0.041 |  |
| Occipital_Mid | -31 | -69 | 31 |  |  |  | 9.69 | <0.001 | 2720 |
|  | -45 | -74 | 13 |  |  |  | 8.38 | 0.001 |  |
| Angular | -52 | -67 | 25 |  |  |  | 7.40 | 0.007 |  |
| undefined | -44 | -15 | -11 |  |  |  | 9.04 | 0.001 | 343 |
| Occipital_Mid | -61 | -46 | -7 |  |  |  | 8.36 | 0.002 | 3468 |
| Temporal_Inf | -54 | -56 | -8 |  |  |  | 7.59 | 0.005 |  |
| Temporal_Mid | -50 | -49 | 10 |  |  |  | 7.56 | 0.006 |  |
| Occipital_Sup | -20 | -64 | 28 |  |  |  | 8.11 | 0.003 | 134 |
| undefined |  |  |  | 43 | -14 | -9 | 7.10 | 0.010 | 77 |
| Occipital_Mid |  |  |  | 51 | -70 | 30 | 6.99 | 0.011 | 269 |
|  |  |  |  | 43 | -73 | 30 | 6.88 | 0.013 |  |
|  |  |  |  | 27 | -75 | 28 | 6.84 | 0.014 | 45 |
|  |  |  |  | 34 | -76 | 40 | 6.34 | 0.027 | 86 |
| ParaHippocampal |  |  |  | 35 | -39 | -5 | 6.55 | 0.020 | 10 |
| Angular |  |  |  | 44 | -53 | 24 | 6.55 | 0.020 | 34 |
| Temporal_Mid |  |  |  | 61 | -42 | -5 | 6.39 | 0.025 | 59 |

**Table S3.**
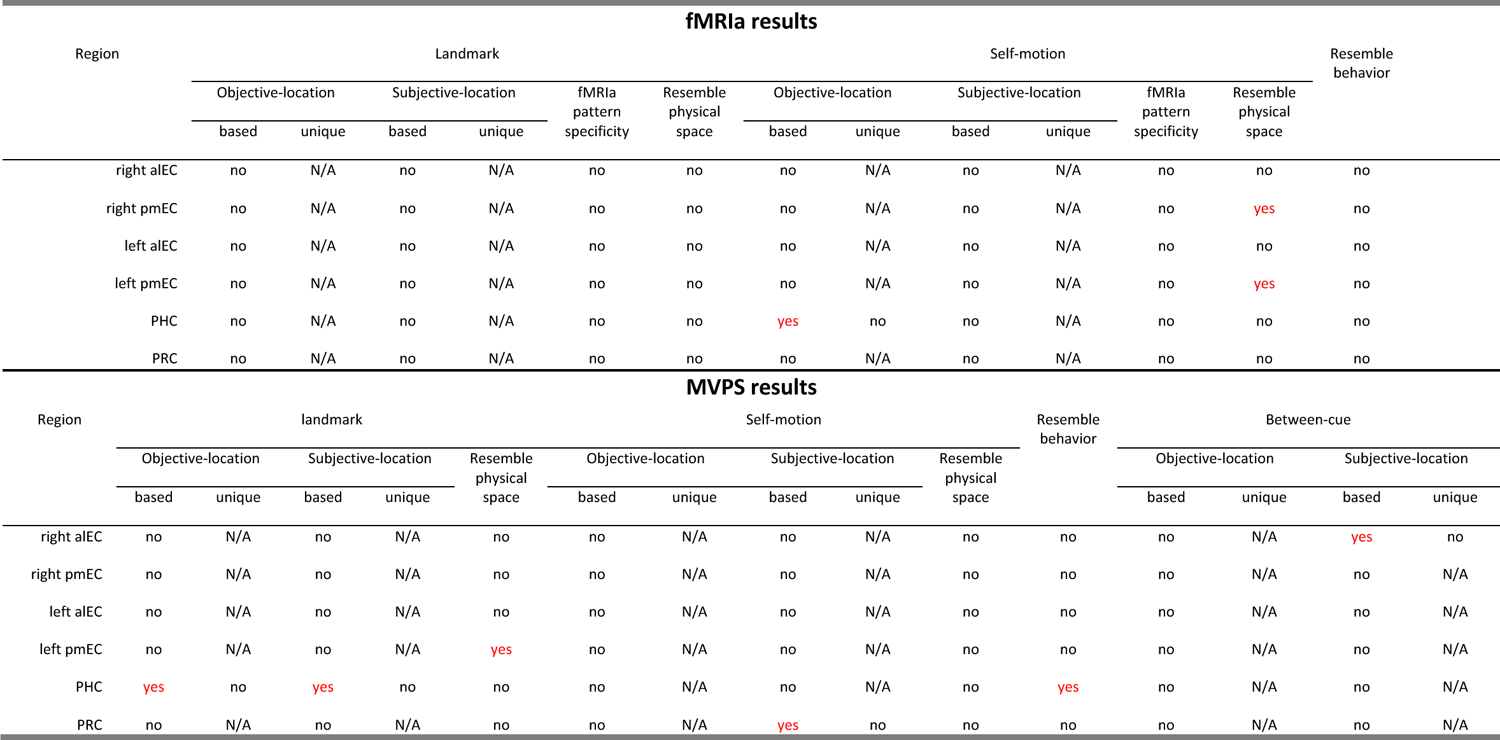
fMRIa and MVPS results of other regions in the medial temporal lobe, related to Figure 3&4&5&6. These areas were analyzed using the same methods as the retrosplenial cortex and hippocampus. Note that we did not attempt to disentangle the unique contributions of objective location and subjective location if the overall fMRIa or MVPS effects were not significant (i.e., N/A). ‘no’ denotes non-significant effect; ‘yes’ denotes significant effect at p < 0.05 (highlighted in red); ‘N/A’ denotes not applicable. alEC: anterior-lateral entorhinal cortex; pmEC: posterior-medial entorhinal cortex; PHC: parahippocampal cortex; PRC: perirhinal cortex.

| fMRIa results |  |  |  |  |  |  |  |  |  |  |  |  |  |
| --- | --- | --- | --- | --- | --- | --- | --- | --- | --- | --- | --- | --- | --- |
| Region | Landmark |  |  |  |  |  | Self-motion |  |  |  |  |  | Resemble behavior |
|  | Objective-location |  | Subjective-location |  | fMRIa pattern specificity | Resemble physical space | Objective-location |  | Subjective-location |  | fMRIa pattern specificity | Resemble physical space |  |
|  | based | unique | based | unique |  |  | based | unique | based | unique |  |  |  |
| right aIEC | no | N/A | no | N/A | no | no | no | N/A | no | N/A | no | no | no |
| right pmEC | no | N/A | no | N/A | no | no | no | N/A | no | N/A | no | yes | no |
| left aIEC | no | N/A | no | N/A | no | no | no | N/A | no | N/A | no | no | no |
| left pmEC | no | N/A | no | N/A | no | no | no | N/A | no | N/A | no | yes | no |
| PHC | no | N/A | no | N/A | no | no | yes | no | no | N/A | no | no | no |
| PRC | no | N/A | no | N/A | no | no | no | N/A | no | N/A | no | no | no |

| MVPS results |  |  |  |  |  |  |  |  |  |  |  |  |  |  |  |
| --- | --- | --- | --- | --- | --- | --- | --- | --- | --- | --- | --- | --- | --- | --- | --- |
| Region | landmark |  |  |  |  | Self-motion |  |  |  |  | Resemble behavior | Between-cue |  |  |  |
|  | Objective-location |  | Subjective-location |  | Resemble physical space | Objective-location |  | Subjective-location |  | Resemble physical space |  | Objective-location |  | Subjective-location |  |
|  | based | unique | based | unique |  | based | unique | based | unique |  |  | based | unique | based | unique |
| right aIEC | no | N/A | no | N/A | no | no | N/A | no | N/A | no | no | no | N/A | yes | no |
| right pmEC | no | N/A | no | N/A | no | no | N/A | no | N/A | no | no | no | N/A | no | N/A |
| left aIEC | no | N/A | no | N/A | no | no | N/A | no | N/A | no | no | no | N/A | no | N/A |
| left pmEC | no | N/A | no | N/A | yes | no | N/A | no | N/A | no | no | no | N/A | no | N/A |
| PHC | yes | no | yes | no | no | no | N/A | no | N/A | no | yes | no | N/A | no | N/A |
| PRC | no | N/A | no | N/A | no | no | N/A | yes | no | no | no | no | N/A | no | N/A |

**Table S4.**
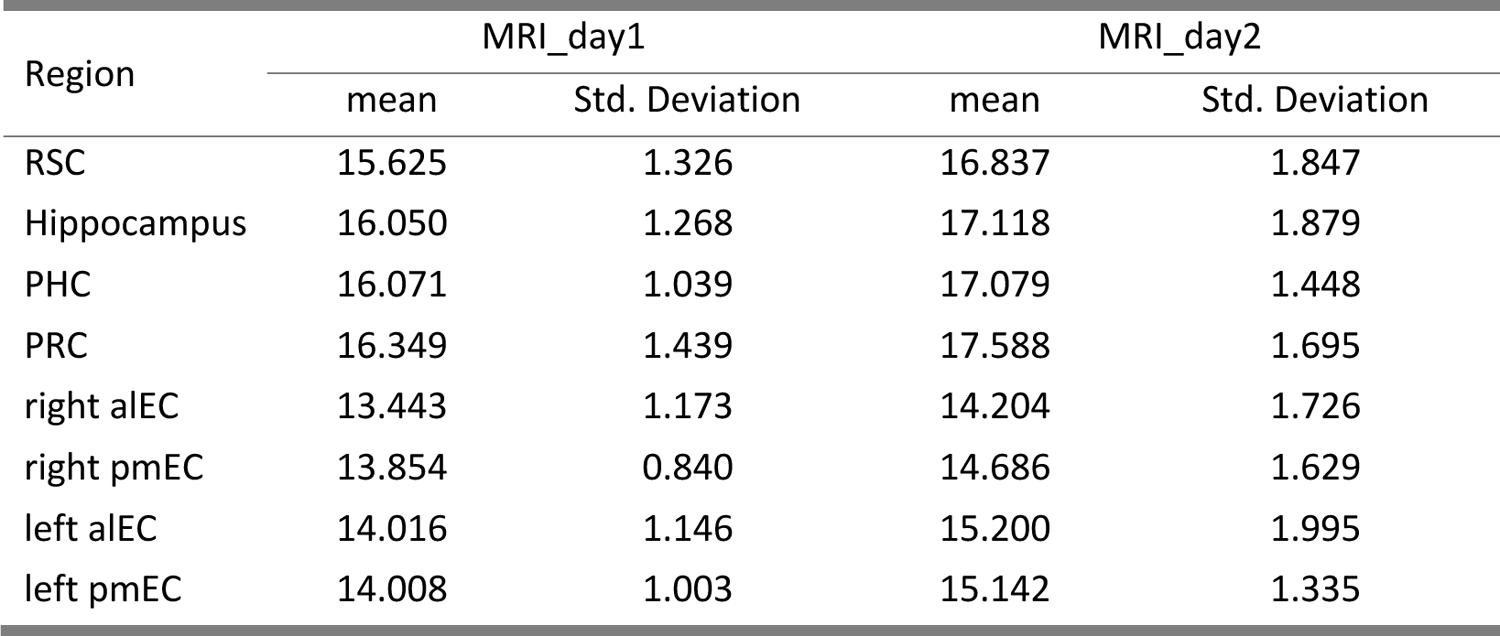
Temporal signal-to-noise ratio (tSNR), related to the discussion section in the main text. tSNR was calculated for each voxel, which was then averaged across all voxels in the brain region. We submitted tSNR to a repeated-measures ANOVA test, with brain region (= 5; the four EC subregions were grouped together), day, and run as independent variables. The main effect of brain region was significant (F(4,76) = 65.432, p < 0.001, η_p_^2^ = 0.775), meaning that the regions differed in tSNR. Post-hoc comparisons with Bonferroni-Holm correction showed that PRC had higher tSNR than RSC, hippocampus, and PHC (ps_corrected_ < 0.001), which in turn had higher tSNR than EC (ps_corrected_ < 0.001). The main effect of day was significant (F(1,19) = 16.422, p < 0.001, η_p_^2^ = 0.464), and there were no significant interaction effects involving day, meaning that for all regions, tSNR significantly improved on the 2^nd^ than the 1^st^ scanning day. The interaction between region and run was significant (F(28,532)=2.634, p < 0.001). Following-up analyses showed that for RSC and PHC, the main effect of run was significant (ps < 0.02), meaning that tSNR decreased linearly across runs (the linear trend of run was significant, RSC, t=4.905, p < 0.001; PHC, t = 4.383, p < 0.001), whereas the main effect of run was non-significant for other regions (ps > 0.07). We also looked more closely at EC by dividing it to four subregions, which were submitted to a repeated-measure ANOVA test, with hemisphere (left vs. right) and entorhinal subregion (alEC vs. pmEC) as independent variables. The main effect of hemisphere was significant (F(1,19) = 39.179, p < 0.001, η_p_^2^ = 0.673), meaning that the left EC had higher tSNR than the right EC. The main effect of subregion was not significant (F(1,19) = 1.503, p = 0.235, η_p_^2^ = 0.073). The interaction between hemisphere and subregion was significant (F(1,19) = 4.560, p = 0.046, η_p_^2^ = 0.194), in that alEC showed greater hemispheric specificity than pmEC. alEC: anterior-lateral entorhinal cortex; pmEC: posterior-medial entorhinal cortex; PHC: parahippocampal cortex; PRC: perirhinal cortex.

| Region | MRI_day1 |  | MRI_day2 |  |
| --- | --- | --- | --- | --- |
|  | mean | Std. Deviation | mean | Std. Deviation |
| RSC | 15.625 | 1.326 | 16.837 | 1.847 |
| Hippocampus | 16.050 | 1.268 | 17.118 | 1.879 |
| PHC | 16.071 | 1.039 | 17.079 | 1.448 |
| PRC | 16.349 | 1.439 | 17.588 | 1.695 |
| right aIEC | 13.443 | 1.173 | 14.204 | 1.726 |
| right pmEC | 13.854 | 0.840 | 14.686 | 1.629 |
| left aIEC | 14.016 | 1.146 | 15.200 | 1.995 |
| left pmEC | 14.008 | 1.003 | 15.142 | 1.335 |

**Table S5.** Behavioral performance over the entire course of experiment, related to Figure 2 and the discussion section of the main text. First, to evaluate the influences of day, we submitted accuracy data to a repeated-measures ANOVA, with day (Pre-scan vs. MRI_day1 vs. MRI_day2) and cue type (landmark vs. self-motion), and run (4 runs) as independent variables. The main effect of day was significant (F(2,38)=13.697 p < 0.001, η_p_^2^ = 0.419). Post-hoc tests showed that the two MRI scanning days did not differ from each other in accuracy (p_holm_ = 0.306), whereas the two scanning days had significantly higher accuracy than the pre-scan day (ps_holm_ = 0.001), indicating that while participants’ performance improved on the first scanning day compared to the pre-scan day, their performance stayed unchanged during the two scanning days. Main effect of cue type was significant (p < 0.001). No other effects were significant (ps > 0.1). These results indicate that the performance improvement mainly occurred between the behavioral training day and the first scanning day. Second, we looked into more details and tested whether behavioral performance changed over time within each day, by submitting behavioral accuracy into repeated-measures ANOVA tests, with cue type (landmark vs. self-motion) and run (4 runs) as independent variables. In the pre-scan day, the main effect of run was significant (F(3,57) = 3.520, p = 0.037, η_p_^2^ = 0.156), and the linear trend of run was significant (t = 2.911, p = 0.005), meaning that behavioral accuracy gradually increased over time. By contrast, the main effect of run was not significant in either the first MRI scanning day (F(3,57) = 0.101, p = 0.959, η_p_^2^ = 0.005; linear trend, t = 0.386, p = 0.701) or the second MRI scanning day (F(3,57) = 1.561, p = 0.209, η_p_^2^ = 0.076; linear trend, t = 0.802, p = 0.426), meaning that behavioral accuracy remained rather stable over time within the day. These results indicated that while there was learning during the first pre-scan training day, no learning occurred during each of the two MRI scanning days.

| Day session |  | landmark |  | self-motion |  |
| --- | --- | --- | --- | --- | --- |
|  |  | Mean | Std. Deviation | Mean | Std. Deviation |
| Pre-scan_day | run1 | 0.766 | 0.137 | 0.644 | 0.169 |
|  | run2 | 0.797 | 0.113 | 0.713 | 0.174 |
|  | run3 | 0.781 | 0.134 | 0.709 | 0.169 |
|  | run4 | 0.829 | 0.116 | 0.734 | 0.146 |
| MRI_day1 | run1 | 0.843 | 0.134 | 0.770 | 0.126 |
|  | run2 | 0.848 | 0.129 | 0.777 | 0.094 |
|  | run3 | 0.840 | 0.143 | 0.792 | 0.123 |
|  | run4 | 0.838 | 0.137 | 0.787 | 0.113 |
| MRI_day2 | run1 | 0.853 | 0.121 | 0.802 | 0.125 |
|  | run2 | 0.848 | 0.123 | 0.762 | 0.153 |
|  | run3 | 0.875 | 0.116 | 0.802 | 0.150 |
|  | run4 | 0.867 | 0.123 | 0.792 | 0.141 |

**Table S6.** ROI-based analyses of navigational success effect in the current study and our previous study (Chen et al., 2019), related to the discussion section in the main text. In the current study, we constructed a GLM, in which the location occupation period of correct trials and incorrect trials were modeled with different regressors. Landmarks and self-motion cues were modeled with different regressors. Because some participants did not make any mistakes in the landmark condition in some runs, scans were concatenated across all the runs in SPM12. Other aspects of the GLM were the same as fMRIa-GLM2, but with no parametric regressors included. For each brain region, mean beta estimate of brain activation was submitted into a repeated measures ANOVA with cue type (landmark vs. self-motion) and correctness (correct vs. incorrect) as independent variables. The results showed that in the current study, only RSC and PHC exhibited significant effects of successful navigation, in that they were more strongly activated in correct trials than in incorrect trials. By contrast, the same analysis in our previous study showed that all the medial temporal lobe areas and RSC significantly contributed to successful navigation. Significant results are highlighted in red. Since there were two more participants in our previous study than in the current study (20 vs. 22 participants), the effect size (i.e., η_p_^2^) is more comparable between the two studies. Results with ROI-specific statistical outliers excluded are in parentheses. RSC: retrosplenial cortex; EC: entorhinal cortex; PHC: parahippocampal cortex; PRC: perirhinal cortex.

| Region | Current study |  |  | Chen et al., 2019 |  |  |
| --- | --- | --- | --- | --- | --- | --- |
| | F | p | $\eta_p^2$ | F | p | $\eta_p^2$ |
| right EC | <0.001<br>(0.195) | 0.982<br>(0.664) | <0.001<br>(0.011) | 9.344<br>(9.01) | 0.006<br>(0.007) | 0.308<br>(0.311) |
| left EC | 3.829<br>(1.461) | 0.065<br>(0.243) | 0.168<br>(0.079) | 6.250 | 0.021 | 0.229 |
| RSC | 17.267<br>(15.512) | 0.001<br>(0.001) | 0.476<br>(0.463) | 20.028<br>(20.196) | <0.001<br>(<0.001) | 0.488<br>(0.502) |
| Hippocampus | 1.375<br>(0.695) | 0.255<br>(0.416) | 0.068<br>(0.037) | 11.886 | 0.002 | 0.361 |
| PHC | 6.282 | 0.021 | 0.248 | 12.309<br>(11.449) | 0.002<br>(0.003) | 0.370<br>(0.364) |
| PRC | 0.823 | 0.376 | 0.775 | 15.304 | < .001 | 0.422 |

